# STREAMING-tag system reveals spatiotemporal relationships between transcriptional regulatory factors and transcriptional activity

**DOI:** 10.1101/2022.01.06.472721

**Authors:** Hiroaki Ohishi, Seiru Shimada, Satoshi Uchino, Jieru Li, Yuko Sato, Manabu Shintani, Hitoshi Owada, Yasuyuki Ohkawa, Alexandros Pertsinidis, Takashi Yamamoto, Hiroshi Kimura, Hiroshi Ochiai

**Affiliations:** Graduate School of Integrated Sciences for Life, Hiroshima University, Higashi-Hiroshima, 739-0046, Japan; School of Life Science and Technology, Tokyo Institute of Technology, Yokohama, 226-8501, Japan; Structural Biology Program, Memorial Sloan Kettering Cancer Center, New York, NY, 10065, USA; Cell Biology Center, Institute of Innovative Research, Tokyo Institute of Technology, Yokohama, 226-8503, Japan; Division of Transcriptomics, Medical Institute of Bioregulation, Kyushu University, Fukuoka, 812-8582, Japan

**Author notes:** Correspondence,; Corresponding authors’ twitter handles: @Cell_Tokyo_Tech, @Hiro_Ochiai_En.

**Keywords:** Transcription, RPB1, BRD4, Mediator, mRNA, RNA polymerase II, live imaging, mintbody, TSS

## Abstract

Transcription is a dynamic process that stochastically switches between the ON and OFF states. To detect the dynamic relationship among protein clusters of RNA polymerase II (RNAPII) and coactivators, gene loci, and transcriptional activity, we inserted an MS2 repeat, a TetO repeat, and inteins with a selection marker just downstream of the transcription start site (TSS). By optimizing the individual elements, we have developed the Spliced TetO REpeAt, MS2 repeat, and INtein sandwiched reporter Gene tag (STREAMING-tag) system. Clusters of RNAPII and BRD4 were observed proximally to the TSS of *Nanog* when the gene was transcribed in mouse embryonic stem cells. In contrast, clusters of MED19 and MED22 Mediator subunits were constitutively located near the TSS. Thus, the STREAMING-tag system revealed the spatiotemporal relationships between transcriptional activity and protein clusters near the gene. This powerful tool is useful for quantitatively understanding dynamic transcriptional regulation in living cells.

## Introduction

In multicellular organisms, a specific gene set is expressed in a particular cell type to support cellular functions. Recent studies involving chromosome conformation capture and its derivatives indicate that promoters and distal enhancers interact with each other to activate the expression of specific genes (Oudelaar and Higgs, 2020). In particular, genomic regions with multiple enhancers are called super enhancers, which contain multiple binding sites for transcription factors involved in cell identity determination (Boija et al., 2018; Hnisz et al., 2013, 2017; Whyte et al., 2013). Transcription factors recruit the transcription machinery and coactivators, including the non-phosphorylated form of RNA polymerase II (RNAPII), a Mediator, and the chromatin regulator bromodomain containing 4 (BRD4) (Cramer, 2019). Since these factors form clusters near the transcriptionally active genes (Cho et al., 2016, 2018; Forero-Quintero et al., 2021; Guo et al., 2019; Li et al., 2019, 2020; Sabari et al., 2018), they may promote the efficient formation of the pre-initiation complex (PIC) and facilitate the transcription initiation of genes. Following the assembly of the complex, initiation is triggered by transcription factor IIH (TFIIH) once its cyclin-dependent kinase 7 (CDK7) subunit phosphorylates the Ser5 in the Tyr1-Ser2-Pro3-Thr4-Ser5-Pro6-Ser7 repeat at the C-terminal domain (CTD) of the RNAPII large subunit RPB1 (Harlen and Churchman, 2017). Subsequently, RNAPII escapes the promoter and is released into productive elongation (Boehning et al., 2018; Cramer, 2019).

Many genes are known to be transcribed non-continuously, switching between an ON state, in which they are continuously transcribed by RNAPII, and an OFF state, in which they are not transcribed. This is called transcriptional bursting, a universal phenomenon observed in many species and cell types, including mouse embryonic stem cells (mESCs) (Larsson et al., 2019; Ochiai et al., 2020; Rodriguez and Larson, 2020; Shah et al., 2018). Although transcription elongation factors are involved in the regulation of transcriptional bursting in a subset of genes, their contribution varies from gene to gene, suggesting that the regulatory mechanism of transcriptional bursting can vary among individual genes (Ochiai et al., 2020). Therefore, to understand the general and gene-specific regulatory mechanisms, further investigation of individual endogenous genes is required.

The MS2 system has often been used as a method to visualize nascent transcripts or transcriptional bursting in living cells (Rodriguez and Larson, 2020; Sato et al., 2020). The focal accumulation of RNAPII and several regulatory complexes has been detected in the vicinity of pluripotency genes in the ON state in mESCs (Cho et al., 2018; Li et al., 2019, 2020). The interaction between enhancers and promoters has also been investigated using MS2 and related systems. The enhancer-gene interaction has shown to be critical in the dynamic regulation of the transcriptional state of certain model reporter genes in *Drosophila* embryos (Chen et al., 2018; Fukaya et al., 2016). However, their interaction was not associated with the transcriptional bursting of the *Sox2* gene in mESCs (Alexander et al., 2019). Since the MS2 system often inhibits translation when inserted at the 5ʹ proximal exon of the gene (Stripecke et al., 1994), the MS2 repeats are usually inserted downstream of the stop codon in the 3ʹ UTR. Since the transcription rate is not always constant (Liu et al., 2021), it is difficult to determine the time delay between transcription initiation and nascent RNA detection. In addition, the intranuclear gene positions in the OFF state cannot be visualized using only the MS2 system. These technical limitations precluded a more detailed analysis of the dynamics of the clusters involving RNAPII and co-factors in relation to the ON/OFF bursting cycles. Therefore, it is demanding to visualize the gene locus during the transcriptional ON and OFF states in living cells.

We previously developed the Real-time Observation of Localization and EXpression (ROLEX) system to visualize the transcriptional activity and intranuclear localization of a specific endogenous gene by the combined use of MS2 and dCas9 systems (Ochiai et al., 2015). However, MS2 repeats were inserted at the 3ʹ UTR in the ROLEX system, and somehow the signal-to-noise ratio of dCas9-mNeonGreen at the MS2 repeat DNA was not very high. Hence, in the present study, we have developed a novel system for the simultaneous quantification of the nuclear localization and transcriptional activity of the gene regions near the TSS, termed as “Spliced TetO REpeAt, MS2 repeat, and INtein sandwiched reporter Gene tag (STREAMING-tag)” system (Fig. 1A).

**Fig. 1.**
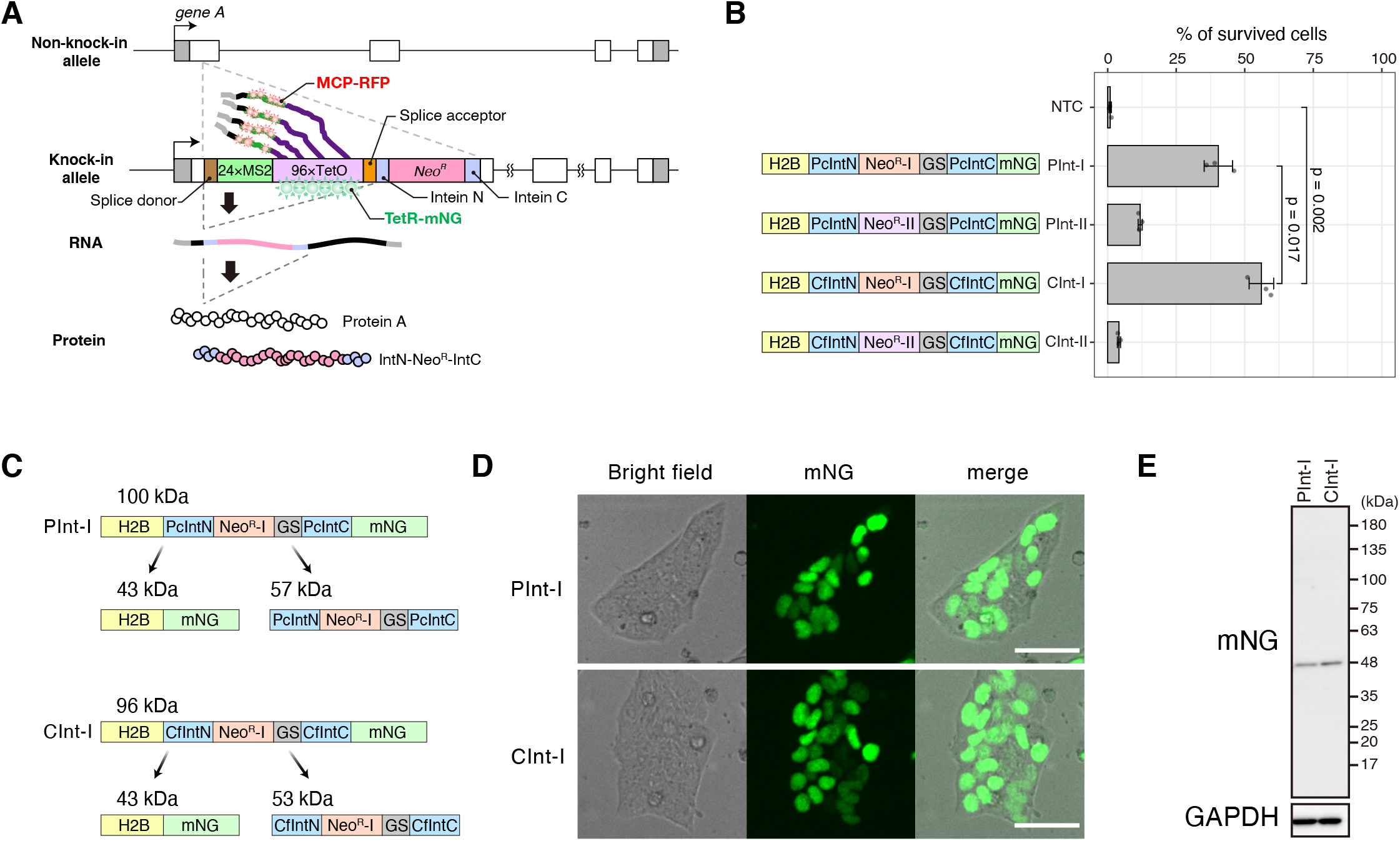
Optimization of selection markers flanked by split inteins. (A) Molecular structure and design of the STREAMING-tag. 24×MS2, 24× Sirius MS2 repeat; 96×TetO, 96× optimized TetO repeat. (B) Expression vectors containing split inteins with NeoR that encodes G418 resistance protein were introduced into cells, and the percentage of cells that survived G418 selection was measured. Data are presented as the means of three biological replicates, and error bars indicate standard deviations. P-values correspond to unpaired, two-sided Student’s t-test. GS, GS linker; NTC, no transfection control. (C) Theoretical structure and size of PInt-I and CInt-I. (D) PInt-I and CInt-I cells established after G418 selection in (B). mNeonGreen (mNG; maximum intensity projections of confocal sections) and bright-field images are shown with their merges. Scale bar, 50 µm. (E) Western blot analysis of PInt-I and CInt -I cells using mNG and GAPDH antibodies. See also Figure S1.

### Design

The STREAMING-tag contains an imaging module comprising MS2 and TetO repeats flanked by a splice donor (SD) and a splice acceptor (SA). The imaging module is followed by a G418-selection module containing *IntN-Neo^R^-IntC*, which encodes an aminoglycoside O-phosphotransferase (APH(3’)), a G418 resistance protein, flanked by split inteins (Fig. 1A). Notably, we have utilized the Sirius MS2 (Ma et al., 2018) and optimized TetO repeats (Tasan et al., 2018) to minimize repetitive sequences. The STREAMING-tag is knocked into a protein-coding region near the TSS. Once the STREAMING-tag is transcribed, the MS2 RNA can be visualized using the MS2 coat protein (MCP) tagged with mScarlet-I, the red fluorescent protein (RFP). The intra-nuclear position of the TetO repeat can be visualized by TetR tagged with mNeonGreen (TetR-mNG). Because the imaging module is flanked by SD and SA and APH(3’) is flanked by split inteins (Ramsden et al., 2011; Stevens et al., 2016), these modules are eliminated by splicing during RNA maturation and protein splicing, respectively. In addition to the amino acids derived from the SD and SA sequences (QG), two amino acids (CF) are added at the N-terminus of C-extein to enhance protein splicing (Stevens et al., 2017), resulting in the insertion of QGCF sequence into the knock-in site of the final protein product. Therefore, the STREAMING-tag-based system is capable of selecting the knocked-in cells using a selection marker, with minimal effect on the target gene function.

## Results

### Optimization of the selection marker flanked by split inteins

First, we optimized the *IntN*-*Neo^R^*-*IntC* gene cassette that encodes APH(3’) flanked by the split inteins. The functionality of type-I aminoglycoside O-phosphotransferase (APH(3’)-I) from *Klebsiella pneumoniae* sandwiched between split inteins from *Penicillium chrysogenum* PRP8 (PcInt) has already been demonstrated (Ramsden et al., 2011). We compared the *Neo^R^*-*I* (encoding APH(3’)-I) and *Neo^R^-II* (encoding APH(3’)-II, which is 35.1% identical to APH(3’)-I; Fig. S1A) sandwiched with either PcInt or a recently reported highly active form of intein Cfa (Stevens et al., 2016). Histone H2B and mNG fusion proteins with different IntN-Neo^R^-IntC modules were introduced into mESCs using the piggyBac system, and the number of G418 resistant cells was examined (Fig. 1B). Neo^R^-I yielded higher resistance than Neo^R^-II, both with PcInt and Cfa (Fig. 1B). Furthermore, Cfa with Neo^R^-I (CInt-I) was more effective than PcInt with Neo^R^-I (PInt-I). Next, we confirmed the nuclear localization and size of mNG in cells transfected with the two constructs conferring G418-resistance (PInt-I and CInt-I) by fluorescence imaging (Fig. 1C, D) and western blotting (Fig. 1C, E). H2B-mNG bands of the anticipated size were observed in both constructs, indicating that the inteins were efficiently excised (Fig. 1E). Therefore, we used the most effective CInt-I for the STREAMING-tag.

### Knock-in of STREAMING-tag into the endogenous *Nanog* gene of mESCs

We knocked-in the STREAMING-tag cassette, consisting of SD, 24× Sirius MS2, 96× optimized TetO, SA, and CInt-I into the TSS-proximal coding region of *Nanog* in mESCs (Fig. 2A S1B-C). Since *Nanog* has multiple transcript variants with distinct TSSs, we selected the knock-in site such that it allows in-frame translation in all the variants; the distances between TSSs and the knock-in site were 311, 314, and 390 bp (Fig. S1D). We generated both single- and double-allele *Nanog*-STREAMING knock-in cell lines (NSt and bNSt, respectively) (Fig. 2B). In the NSt cells, a six-nucleotide deletion was introduced in the non-knocked-in allele (Fig. S1E).

**Fig 2.**
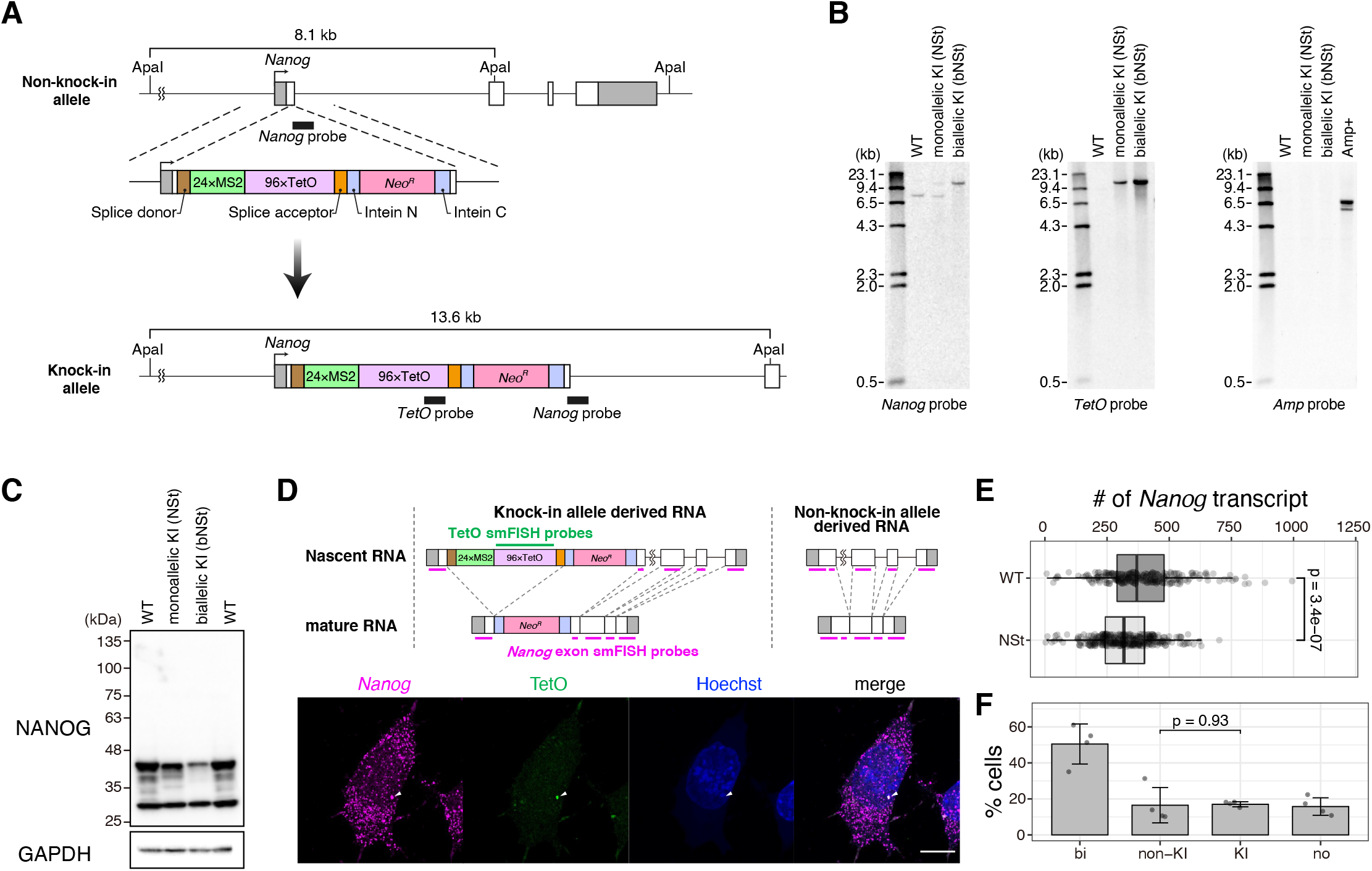
STREAMING-tag knock-in into endogenous Nanog in mouse embryonic stem cells (mESCs). (A) Mouse Nanog gene structure after STREAMING-tag knock-in. (B) Southern blot analysis of single- and double-allele Nanog-STREAMING knock-in (NSt and bNSt, respectively), and wild-type (WT) cells. (C) Western blot analysis of NSt, bNSt, and WT cells. (D–F) Single-molecule fluorescence in situ hybridization (smFISH) analysis of NSt and WT cells. (D) Upper panel shows the location of the smFISH probes, and lower panels show example images of NSt cells. Arrowheads indicate Nanog transcriptional spots. Scale bar, 10 µm. (E) Distribution of Nanog mRNA counts in WT and NSt cells. WT, N = 214 cells; NSt, N = 278 cells. P-values were determined using Wilcoxon rank sum test. (F) Percentage of cells showing bright Nanog RNA-FISH signals at the transcription sites in NSt cells. The percentages of cells with RNA-FISH signals at only transcription sites for the non-knock-in allele (non-KI), at only transcription sites for the knock-in allele (KI), at transcription sites for both alleles (bi), and not at transcription sites (no) are plotted [mean ± SD; N = 4 biological replicates (data represented as dots); >80 cells in a single experiment]. P-values correspond to unpaired, two-sided Student’s t-test. See also Figures S1 and S2.

Western blotting analysis showed that NANOG protein levels were slightly and substantially lower in NSt and bNSt cells, respectively, than that in the parental cells (Fig. 2C). Although the *Nanog* gene is important for the maintenance of pluripotency, no obvious abnormalities were observed in the cell morphology of either NSt or bNSt cells. This suggests that even though the level of the NANOG protein decreased with the four amino acid insertion, the protein function may have remained preserved.

To examine the expression of the *Nanog* mRNA and the splicing of the MS2*-*TetO cassette, single-molecule fluorescence *in situ* hybridization (smFISH) was performed on the NSt and WT cells using probes designed against the *Nanog* exon region (Ochiai et al., 2014) and the TetO repeat region (Fig. 2D). *Nanog* smFISH signals were scattered throughout the cytoplasm, with particularly strong fluorescence spots in the cell nucleus, in both the cell types (Fig. 2D, S1F) (Ochiai et al., 2014). smFISH signals of TetO were negligible in the cytoplasm, while a strong fluorescent spot was observed in the nucleus. Since the nuclear TetO smFISH spot co-localized with the *Nanog* fluorescent spot (Fig. 2D), and because introns are rapidly degraded after splicing (Ooi et al., 2001), this spot putatively represents the nascent transcripts before splicing. Quantification of *Nanog* smFISH spots in NSt and WT cells revealed that the number of *Nanog* mRNA in NSt cells was considerably reduced (18% reduction) compared to that in WT cells (Fig. 2E). Furthermore, we analyzed the transcriptional efficiency of the knock-in and non-knock-in alleles in NSt cells and found no significant difference (Fig. 2F). The transcriptional efficiencies of cells showing a single nuclear *Nanog* transcriptional spot, with or without a TetO smFISH spot were 17.0±1.4% and 16.5±9.8%, respectively, whereas that of the cells showing two nuclear *Nanog* spots was 50.5±11.1%. This suggests that STREAMING-tag knock-in does not have a significant effect on transcription initiation and pause-release. Considering the insertion of a significantly long cassette (5.5 kb) in the 7 kb-long *Nanog* gene, the decreased expression of the knock-in allele observed via western blotting and smFISH may be explained by the additional time required for elongation and splicing (Castillo-Davis et al., 2002; Marais et al., 2005; Swinburne and Silver, 2008).

We also established mESCs harboring a STREAMING-tag knock-in at *Sox2*, encoding the pluripotency transcription factor SOX2 and *Usp5*, encoding ubiquitin-specific peptidase 5 (Fig. S2). Western blot analysis showed a moderate decrease in protein expression in monoallelic knock-in cell lines. While *Sox2* (2 kb) is shorter than *Nanog*, *Usp5* is much longer (14.4 kb) than it, and both of their expression levels were affected. Therefore, STREAMING-tag insertion may affect the elongation and splicing of the target gene independent of its length and without influencing transcription initiation and pause-release.

### TetR(W43F) is more suitable than TetR(WT) for DNA labeling

To visualize the location and transcription of the STREAMING-tag in the knock-in allele, we expressed wild-type TetR (TetR(WT))-mNG and MCP-RFP in the NSt cell line (Fig. 3A). In the cells where TetR(WT)-mNG was highly expressed, a single mNG spot was clearly detected in the nucleus, whereas MCP-RFP spot was almost undetectable. In contrast, in the cells showing low expression of TetR(WT)-mNG, the MCP-RFP spots were detectable. This suggests that the transcription of the STREAMING-tag was blocked due to the strong DNA binding of TetR(WT). Therefore, we introduced a mutation in TetR to reduce its DNA-binding affinity without affecting the sequence specificity (Baumeister et al., 1992) (Fig. 3B, C). Among the five mutants tested, each with a single amino acid substitution, TetR(W43F) labeling resulted in detection of both TetR-mNG and MCP-RFP spots in the highest percentage of cells (Fig. 3B, D). To examine the effect of this mutation on the binding property of TetR, we performed fluorescence recovery after photobleaching (FRAP) analysis (Fig. 3E, F). The obtained FRAP curve could be well fitted to a single exponential with the baseline (see STAR Methods). The mobile fraction was not significantly different between TetR(WT) and TetR(W43F) (53.3±11.0 % and 49.2±1.4 % for TetR(WT) and TetR(W43F), respectively). In contrast, the recovery time constant of TetR(W43F) was significantly smaller than that of TetR(WT) (107.4±34.0 s and 26.3±2.8 s for TetR(WT) and TetR(W43F), respectively; *p* = 0.0181, pairwise *t*-test). This indicates that TetR(W43F) has a faster dissociation rate than TetR(WT), resulting in a reduced inhibitory effect on transcription (Fig. 3E, F).

**Fig. 3.**
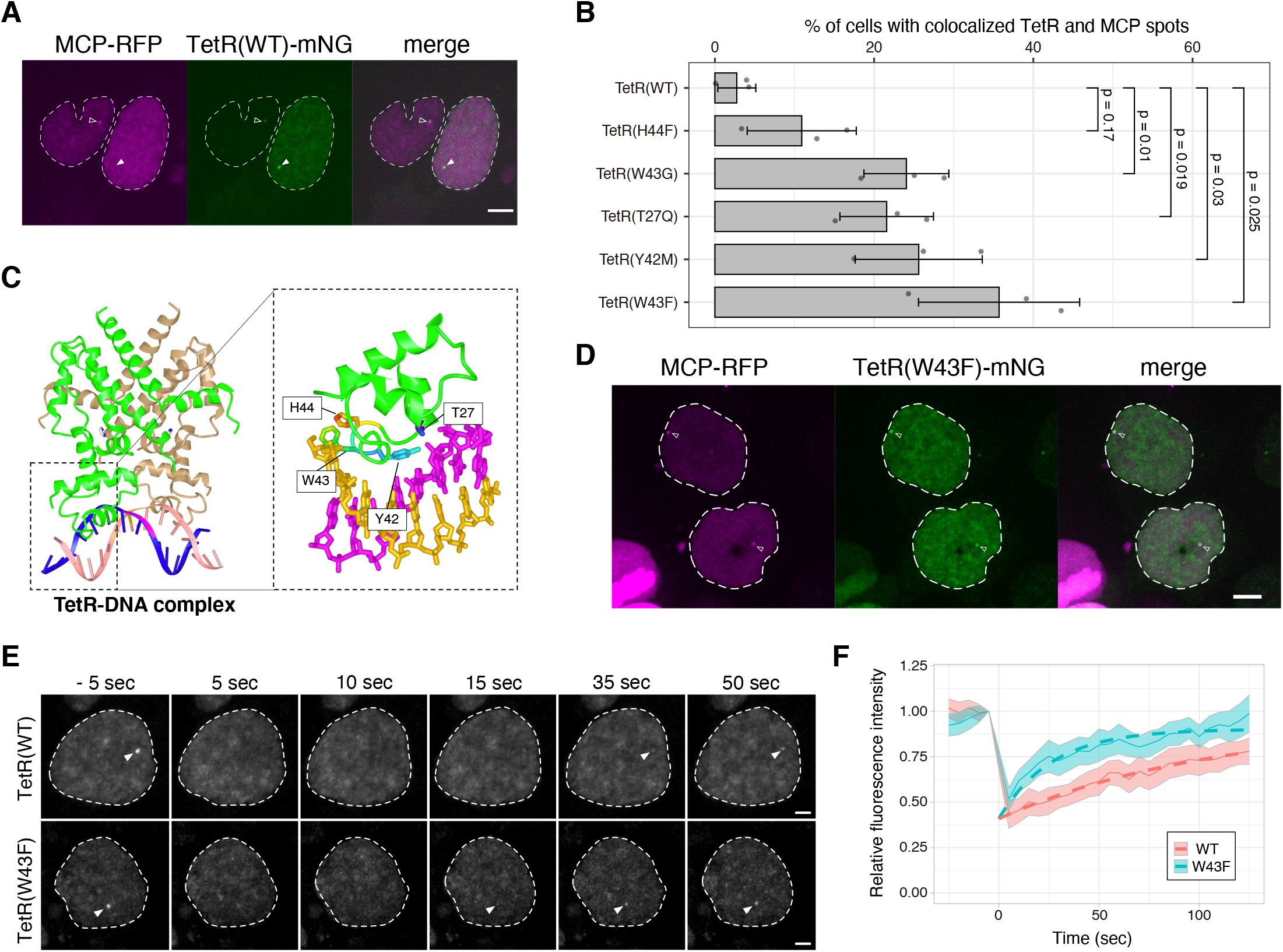
TetR(W43F) is more suitable than TetR(WT) for DNA labeling. (A) Nanog STREAMING-tag knock-in (NSt) cells transiently expressing TetR(WT)-mNG and MCP-RFP. In cells with high TetR(WT)-mNG expression (right), MCP spots that overlapped with TetR(WT) spots were rarely observed (closed arrowhead). In cells with low TetR(WT)-mNG expression (left), MCP spots tended to overlap with TetR(WT) spots (open arrowhead). Dashed line indicates the cell nucleus. Scale bar, 5 µm. (B) Percentage of cells in which TetR and MCP spots were simultaneously visible and colocalized. Mean values with SD of three biological replicates (>30 cells) are shown. P-values correspond to unpaired, two-sided Student’s t-test. (C) 3D structure of TetR (PDB 1QPI) and mutation site location. (D) Images of NSt cells co-transfected with TetR(W43F)-mNG and MCP-RFP. Open arrowheads indicate TetR spots with MCP spots. Dashed line indicates the cell nucleus. Scale bar, 5 µm. (E and F) FRAP assay in NSt cells expressing TetR(WT)-mNG or TetR(W43F)-mNG. (E) FRAP images. TetR-mNG spots were transiently bleached, and the recovery of spots was observed over time. Dashed lines and arrowheads indicate cell nuclei and TetR-mNG spots, respectively. Scale bar, 2 µm. (F) FRAP curves. Relative fluorescence intensities of TetR(WT)- and TetR(W43F)-mNG after photobleaching. Means ± 95% CI (WT, N = 24; W43F, N = 23) are shown as the fitted curves (bold dashed lines) with single exponential fitting with baseline (see STAR Methods in detail). See also Figure S3.

To confirm whether the TetR(W43F)-mNG and MCP-RFP spots observed using the STREAMING-tag system in the living NSt cells truly represent the *Nanog* locus, we performed DNA-FISH (Fig. S3). NSt cells expressing TetR(W43F) and MCP-RFP were fixed and fluorescence images were acquired. Subsequently, DNA-FISH was performed on the same cells using the *Nanog* probe, which were imaged to detect DNA-FISH signals that were compared with protein fluorescence. The fluorescent spots of TetR(W43F)-mNG and MCP-RFP showed significant overlap with one of the DNA-FISH spots (Fig. S3), indicating that the knock-in of the STREAMING-tag can be used to visualize specific gene regions in living cells. TetR(W43F) is hereafter referred to as mutant TetR (mTetR).

### Verification of the versatility of STREAMING-tag

In the ROLEX system, we knocked-in the MS2 repeat immediately downstream of the *Nanog* stop codon to visualize the transcription with MCP-RFP and the MS2 DNA repeat with dCas9-mNG (Ochiai et al., 2015). Compared to the data obtained from dCas9-mNG in the ROLEX system, the ratio of the mean signal intensity of mTetR-mNG spots to the standard deviation of the nuclear area (signal-to-noise ratio; SNR, see STAR Methods) in the STREAMING-tag system was significantly higher, indicating improved visualization of the gene locus in the STREAMING-tag system (Fig. 4A).

**Fig. 4.**
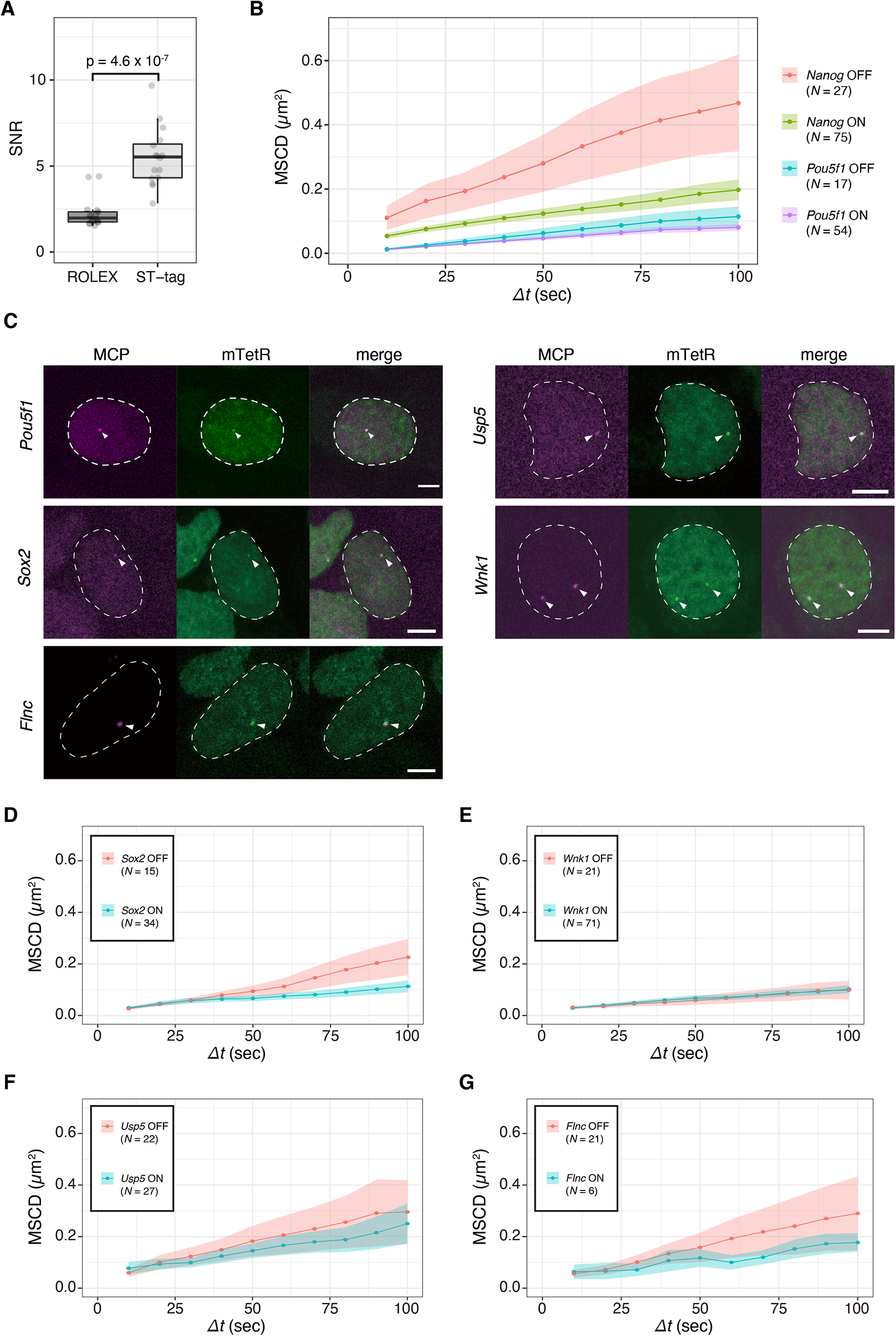
Verification of versatility of the STREAMING-tag. (A) Comparison of the signal-to-noise ratio (SNR) of DNA labeling spots between the ROLEX and STREAMING-tag systems. N = 16 cells. P-value was determined using Wilcoxon rank sum test. (B) Mean square change in distance (MSCD) of mTetR spots with respect to the center of the nucleus in NSt-NLS-SNAP cells for Nanog and PSt-NLS-SNAP cells for Pou5f1. The data are classified into the ON and OFF states. Means with standard error of the mean (SEM) are shown. N, number of cells analyzed. (C) Imaging of mTetR and MCP spots in knock-in cells with the STREAMING-tag into Pou5f1, Sox2, Flnc, Usp5, and Wnk1. Dashed lines and arrowheads indicate cell nuclei and mTetR/MCP spots, respectively. Scale bar, 5 µm. In Wnk1 STREAMING-tag knock-in cells, two spots were observed because the STREAMING-tag was knocked-in into both alleles. (D–G) MSCD of mTetR spots with respect to the center of the nucleus in SSt-NLS-SNAP for Sox2 (D), WSt-NLS-SNAP for Wnk1 (E), USt-NLS-SNAP for Usp5 (F), and FSt-NLS-SNAP cells for Flnc (G). The data are classified into the ON and OFF states. Means with SEM are shown. N, number of cells analyzed. See also Figure S4.

Using the ROLEX system, we reported that the mobility of the *Nanog* locus in mESCs significantly increases in the OFF state compared to that in the ON state, whereas that of *Pou5f1* is independent of its transcriptional state (Ochiai et al., 2015). In the present study, we tested whether similar behaviors are observed in the STREAMING-tag system. The *Nanog* locus showed higher mobility in the OFF state than that in the ON state (Fig. 4B, Movie S1). In contrast, the STREAMING-tag knocked-in the *Pou5f1* locus (Fig. 4C, S4) showed negligible difference in mobility between the ON and OFF states. These results were in accordance with previous data (Ochiai et al., 2015), suggesting that the dynamic behavior of gene loci in living cells can be quantified using the STREAMING-tag system.

To confirm the versatility of the STREAMING-tag, we further knocked-in the STREAMING-tag into other genes, including *Wnk1* and *Flnc* (Fig. S4). We also expressed mTetR-mNG and MCP-RFP in *Sox2* and *Usp5* STREAMING-tag knock-in cells and the transcription and gene loci were clearly observed in all the cell lines (Fig. 4C). Furthermore, in these cells, we observed considerable differences in mobility between the transcriptional ON and OFF states for *Sox2*, but not for *Wnk1*, *Usp5*, and *Flnc* (Fig. 4D-G). These results suggest that STREAMING-tag can be used to track the movement of various genes.

### STREAMING-tag system enables quantification of transcriptional dynamics near TSS

In cells with STREAMING-tag knocked-in near TSSs, the duration between transcription initiation, followed by pause-release and productive elongation, and the appearance of the MCP-tagged nascent RNA spot is expected to be shorter than in cells with the MS2 repeat sequence knocked-in downstream of the stop codon. We compared the time points of MCP spot appearance during transcription restoration in the NM-G cell line (Ochiai et al., 2014) harboring the MS2 repeat knocked-in immediately downstream of the *Nanog* stop codon and in the NSt-GR cell line. We treated the cells with 5,6-dichloro-1-β-d-ribofuranosylbenzimidazole (DRB), a reversible cyclin-dependent kinase 9 (CDK9) inhibitor (Bensaude, 2014), for 90 min, removed it by washing, and recorded the time of the appearance of the transcriptional spot (Fig. 5A, B). As the *Nanog* gene displays both ON and OFF states, the activation of *Nanog* transcription is not anticipated to occur in all cells after DRB release. Therefore, we compared the respective timings of the waves of transcriptional spot appearance. The first wave of cumulative density function (CDF) appeared at 2-3 min in NSt-GR cells and 8-9 min in NM-G cells (Fig. 5C-F). The different timings are consistent with the respective positions of the MS2 repeats in *Nanog* in these cells. Thus, STREAMING-tag knocked-in near TSSs allows the quantification of transcriptional activity immediately after transcription initiation, pause-release, and productive elongation.

**Fig. 5.**
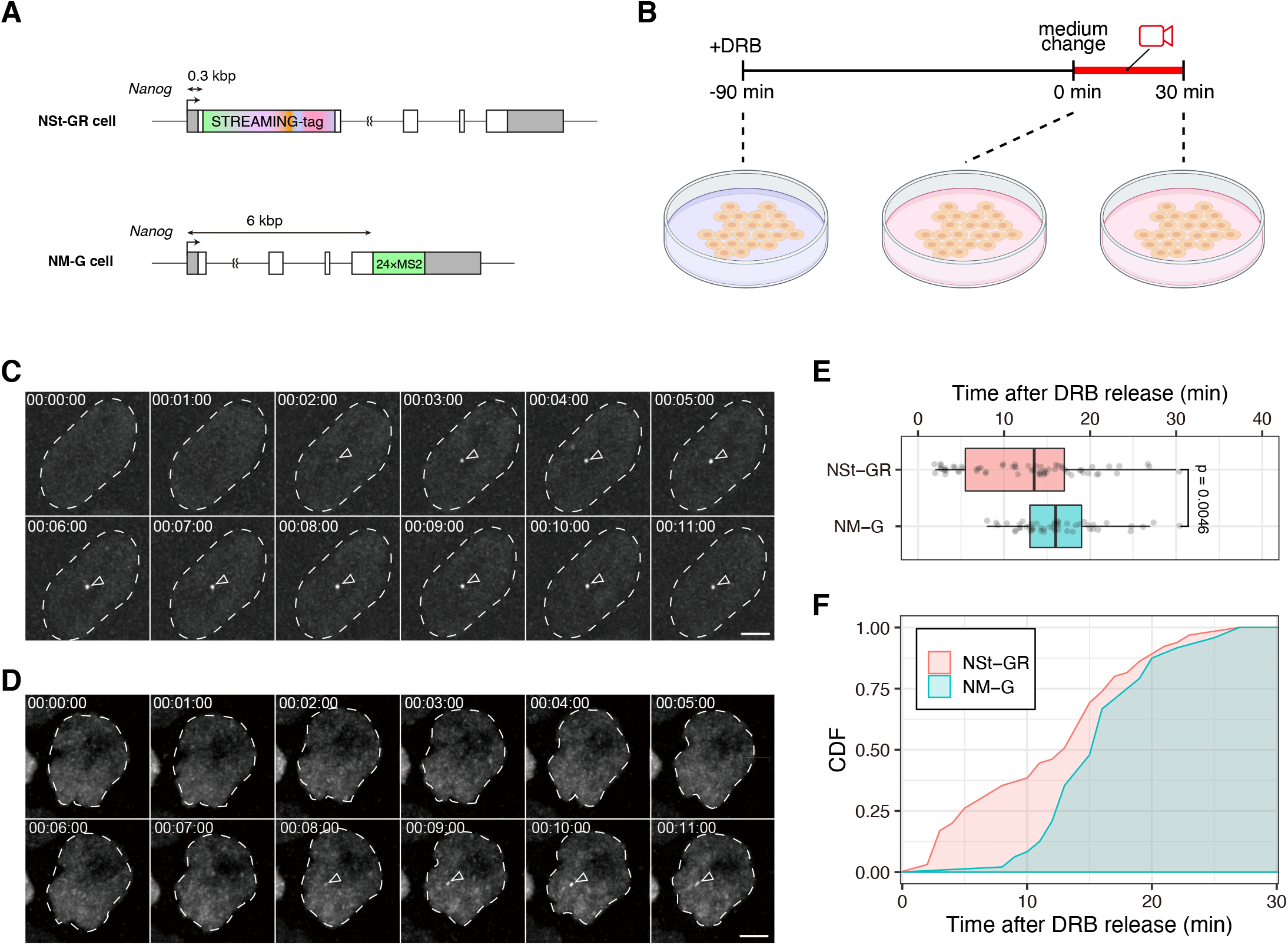
STREAMING-tag system enables quantification of transcriptional dynamics near transcription start site (TSS). (A) Genomic structure of NSt cell line moderately expressing mTetR-mNG and MCP-RFP (NSt-GR), and a cell line in which MS2 repeats were biallelically knocked-in immediately downstream of the Nanog stop codon, moderately expressing MCP-mNG (NM-G). (B) Schematic of 5,6-dichloro-1-β-d-ribofuranosylbenzimidazole (DRB) releasing assay. Cells were treated with DRB for 90 min, washed, and then live-imaged at 1 min intervals for 30 min. (C, D) Live imaging of NSt-GR (C) and NM-G (D) cells after DRB release. Dashed lines indicate cell nuclei. Arrowheads indicate transcription spots. Scale bar, 5 µm. (E) Distribution of the time when the first transcription spot was observed in NSt-GR and NM-G (NSt-GR, N = 66; NM-G, N = 49). Boxes indicate inter-quartile range (IQR; 25–75% intervals) and the median line; whiskers indicate 1.5-fold of the IQR. P-values were determined using Wilcoxon rank sum test. (F) Cumulative density function (CDF) of the data in (E).

### RPB1 and BRD4 form clusters in proximity to *Nanog* only in the ON state in mESCs

We have previously reported that RPB1 and BRD4 form clusters in the vicinity of *Nanog* and *Pou5f1* during the ON state using cell lines in which MS2 repeats were inserted immediately downstream of the stop codon (Li et al., 2019, 2020). However, it was unclear whether these clusters are also formed in the OFF state (Fig. 6A). Therefore, we established several cell lines in which SNAPtag was knocked into *Rpb1*, *Brd4*, *Med19*, and *Med22* in NSt-GR cells and measured the 2D distance between mTetR spots and the nearest transcriptional regulatory factor (RF) clusters in the ON and OFF states (Fig. 6B, C, Fig. S5A-C, Table S1). The distances between the mTetR spot and the nearest RPB1 or BRD4 cluster were significantly shorter in the ON state than in the OFF state (median ∼250 nm and ∼500 nm, respectively). However, the distance between mTetR and the nearest MED19 or MED22 cluster was similar to those between mTetR and RPB1 and BRD4 clusters in the ON state, regardless of their transcription state (Fig. 6 C, D; median < 350 nm).

**Fig. 6.**
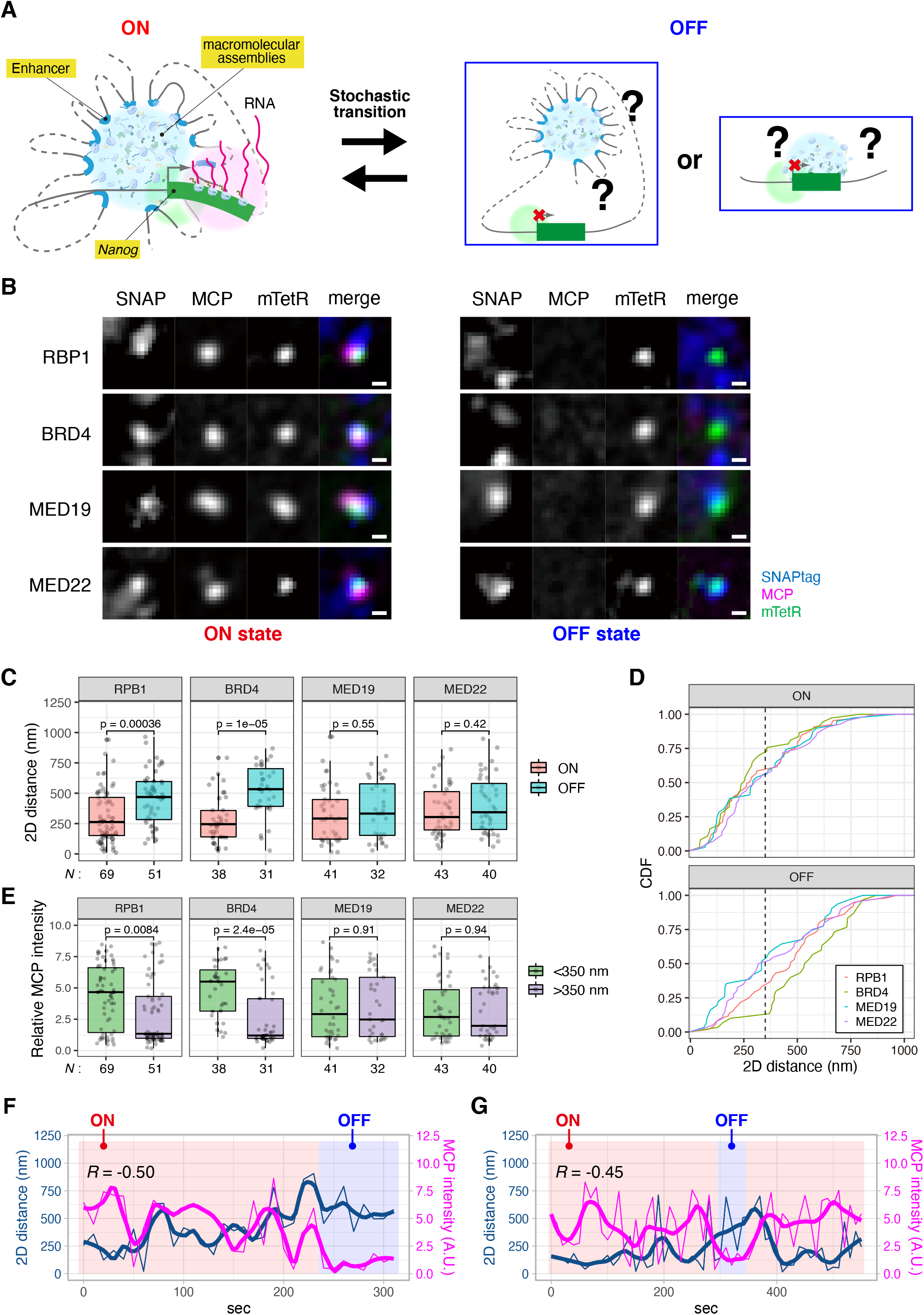
RPB1 and BRD4 form clusters in proximity to Nanog only in the ON state in mESCs. (A) Hypothetical relationship between the transcriptional state and higher-order genomic structures of Nanog. (B) Relationship between MCP, mTetR, and SNAP-tagged regulatory factors (RFs). Using NSt-GR cells as a parental cell line, SNAPtag was knocked-in into RF genes. Images with (ON state; left) and without MCP (OFF state; right) are shown. Scale bar, 500 nm. (C) 2D distances between mTetR spots and the nearest SNAPtag clusters in the ON and OFF states. N, number of cells analyzed. P-values were determined using Wilcoxon rank sum test. (D) Cumulative density function (CDF) of the data in (C). Black dashed line indicates 350 nm. (E) Distributions of MCP fluorescence intensities at the mTetR spot with the 2D distance between the mTetR spot and nearest SNAPtag cluster at <350 and >350 nm. N, number of cells analyzed. (F and G) Time-lapse analysis of MCP, mTetR, and SNAPtag-RFs (F, RPB1; and G, BRD4). 2D distance between mTetR and RF clusters (blue), and MCP fluorescence intensity (magenta) is shown. The thin lines indicate the experimental data, and the thick lines indicate the locally estimated scatterplot smoothing (LOESS) regression with the degree of smoothing α = 0.2, drawn using R ggplot2, to visualize the gradual dynamic change in noisy data. Spearman’s rank correlation coefficient (R) of smoothed lines is shown. See also Figure S5.

Next, we analyzed the correlation between MCP spot fluorescence intensity, as a proxy of transcriptional activity, and the distance between mTetR and the nearest RF clusters. As the median distances of the mTetR-RF clusters are below 350 nm (Fig. 6D) in the ON state, we set a threshold of 350 nm to categorize the distances as short or long. The MCP fluorescence intensity was higher when an RPB1 or BRD4 cluster was within 350 nm of the mTetR spot, while no such systematic correlation was observed in case of MED19 and MED22 (Fig. 6E). These data suggest that RPB1 and BRD4 clusters are not formed near *Nanog* particularly in the OFF state, whereas MED19 and MED22 clusters are formed in the vicinity of *Nanog* independent of their transcriptional state.

### Dynamics of mTetR-RF distances and MCP intensities

To analyze the relationship between the MCP fluorescent intensity (transcriptional activity) and mTetR-RF cluster distance, we performed the live imaging of MCP, mTetR, and RF (RPB1 and BRD4) clusters at 10 s intervals for 10 min (Fig. 6F-G). For this analysis, we acquired five z-sections per stack spanning 0.8 µm (z-step = 200 nm) to minimize the time intervals using minimum laser power for long-term imaging; however, spot tracking was rarely achieved because of the z-direction movements of the spots. Among the spots that were tracked for more than 5 min, anti-correlation trends were sometimes observed between the MCP intensity and the mTetR-RF cluster distance (Fig. 6F-G). These results indicate that the STREAMING-tag system can be used to simultaneously quantify the distance between RF clusters and an endogenous gene locus and monitor the transcriptional activity of the gene over time. In addition, it was inferred that these relationships are highly dynamic in living cells.

### RNAPII Ser5ph and Ser2ph mintbodies form foci in proximity to *Nanog* only in the ON state in mESCs

To investigate which form of RNAPII is associated with *Nanog* during the ON state, we used genetically encoded modification-specific intracellular antibodies (mintbodies), which consist of a single chain variable fragment (scFv) of a specific antibody and a fluorescent protein (Sato et al., 2013; Tjalsma et al., 2021). In addition to the previously established RNAPII Ser2ph-specific mintbody (Uchino et al., 2021), we generated an RNAPII Ser5ph-specific mintbody (Fig. S6, see STAR Methods). We cloned scFv from mouse hybridoma cells producing the RNAPII Ser5ph-specific antibody (Fig. S6A) and introduced amino acid substitutions in the framework region to improve the stability (Fig. S6B-D). We then confirmed its specificity using enzyme-linked immunosorbent assay (ELISA) (Fig. S6E-F) and inhibitor treatments (Fig. S6G-J).

NSt-derived cells expressing mTetR-mNG, MCP-RFP, and mintbody-SNAPtag were established and imaged following the same procedure as RF cluster imaging. For both Ser2ph and Ser5ph mintbodies, enriched foci were observed close to mTetR and MCP in the ON state (Fig. 7A). We then quantified the distance between the mTetR spots and the nearest mintbody foci (Fig. 7B, C, Fig. S5A-C). In case of both RNAPII Ser5ph and Ser2ph, the distance between mTetR and the nearest mintbody foci was significantly shorter in the ON state than that in the OFF state, as observed for RPB1. In addition, the distance from mTetR to the nearest RNAPII Ser5ph foci was similar to that of RPB1 but slightly shorter than that observed for RNAPII Ser2ph foci (Fig. 7B, C). This suggests that the distance of different classes of RNAPII foci from mTetR spot is the following order: the RNAPII Ser5ph foci, the RPB1 cluster potentially including the non-phosphorylated (pre-initiating) or phosphorylated (elongating)-form of RNAPII, and the RNAPII Ser2ph foci. This implies that the STREAMING-tag system can visualize the TSS-proximal region of *Nanog*.

**Fig. 7.**
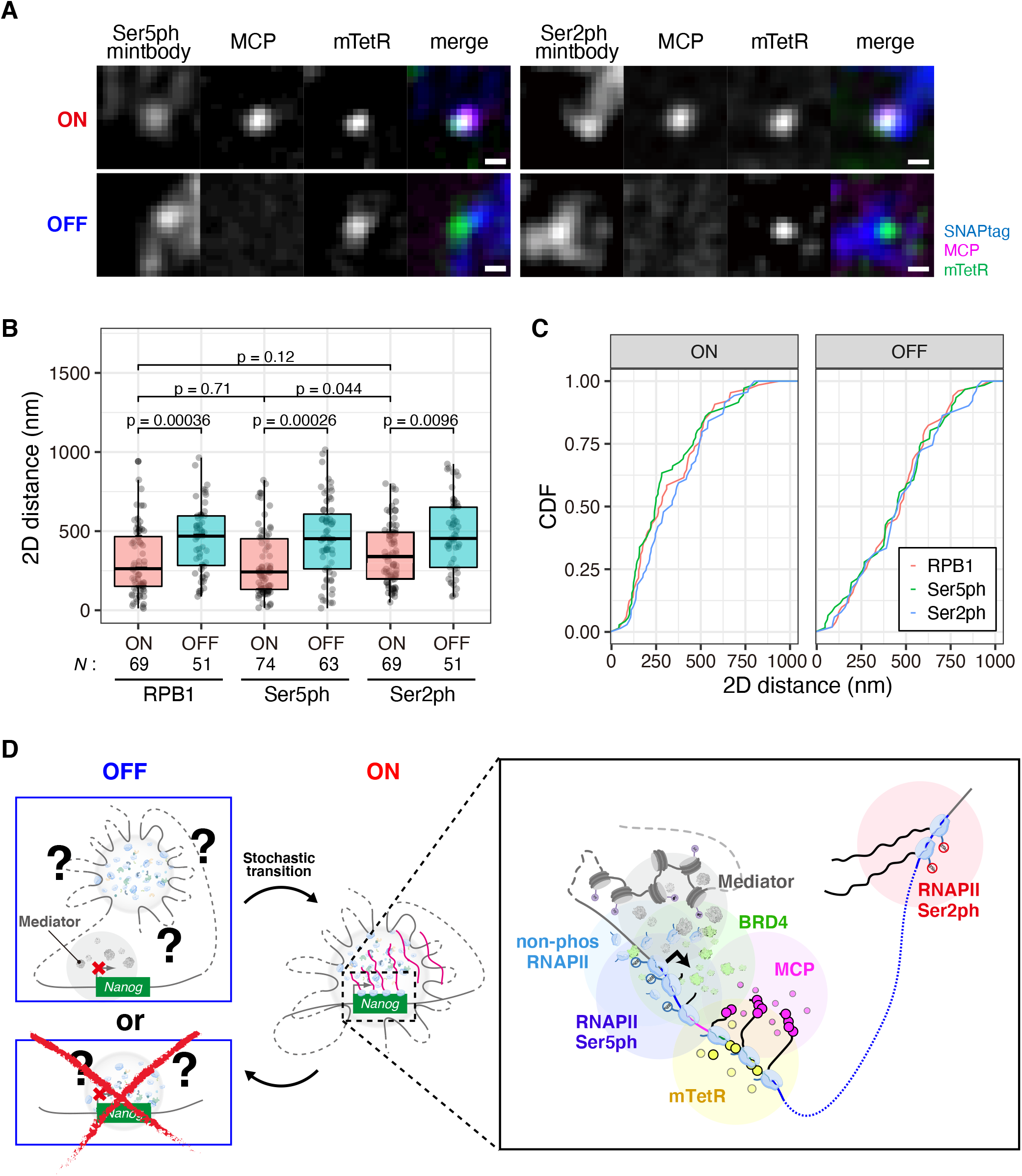
RNAPII Ser5ph and Ser2ph form clusters in proximity to Nanog only in the ON state in mouse embryonic stem cells (mESCs). (A) Images of RNAPII Ser5ph- and Ser2ph-mintbodies with MCP and mTetR. Cell lines expressing RNAPII Ser5ph and RNAPII Ser2ph mintbody-SNAPtag were established using NSt-GR as a parental cell line. Scale bar, 500 nm. (B) 2D distance between mTetR spots and the nearest SNAPtag clusters in the ON and OFF states. RPB1 data are the same as in Fig. 6C, represented for ease of comparison. N, number of cells analyzed. P-values were determined using Wilcoxon rank sum test. (C) Cumulative density function (CDF) of the data in (B). (D) Model of relationship among transcriptional activity, RF clusters, and different forms of phospho-RNAPII clusters in Nanog in mESCs. See also Figures S5 and S6.

## Discussion

Transcription is a dynamic process that is switched stochastically between the ON and OFF states (Rodriguez and Larson, 2020). We previously showed that transcriptional bursting is controlled by multiple factors, depending on the individual genes (Ochiai et al., 2020). Therefore, it is important to determine the bursting kinetics of individual genes, for which live imaging is one of the best approaches. In this study, we demonstrated that the STREAMING-tag knocked into the TSS-proximal coding region allows the simultaneous determination of nuclear localization and transcriptional state of an endogenous gene without significantly affecting gene function.

### Characteristics and advantages of the STREAMING-tag system for understanding regulatory mechanisms of transcriptional bursting

Because the MS2 system can inhibit protein translation, MS2 repeats are usually inserted immediately after the stop codon of endogenous genes (Stripecke et al., 1994). However, in this case, we anticipated a time lag between the actual initiation of productive elongation by RNAPII from the TSS and the fluorescent spot detection by the MS2 system. Since the transcription speed is not constant (Liu et al., 2021), it is challenging to estimate the precise time points of transcription initiation and/or pause-release, which are relevant to the switching of ON and OFF states (Bartman et al., 2018), before the fluorescent spot is observed. In contrast to the standard MS2 system, the STREAMING-tag system can be knocked into the TSS-proximal coding region of various genes, such that the gene regions and transcripts can be simultaneously visualized. This TSS-proximal insertion enables the detection of transcripts immediately after the onset of processive transcription.

We compared the time of transcriptional spot recovery following DRB release using the cells carrying the STREAMING-tag, and MS2 repeats were knocked into the regions proximal to TSS (+0.3 kb) and downstream of the stop-codon (+6 kb), respectively, in the *Nanog* gene. The recovery time was approximately 6 min faster in the STREAMING-tag knock-in cells than in the MS2 repeat knock-in cells. The distance between the STREAMING-tag and MS2 repeat sites was 5.7 kb, and the transcription elongation rate was estimated to be approximately 1 kb/min, which lies within the characteristic range of transcription elongation speed (0.5-4 kb/min) (Jonkers et al., 2014). Thus, the STREAMING-tag knock-in near the TSS is potentially useful for observing the transcriptional state immediately after the onset of productive elongation and consequently in understanding the mechanism of transcriptional regulation.

Furthermore, the STREAMING-tag system permits the comparison of the localization of RFs involved in transcriptional bursting to the TSS-proximal region (Fig. 7D). In a previous study, the RPB1 cluster was observed to be closer to the MCP transcriptional spot than to the BRD4 cluster in a cell line wherein the MS2 repeats were inserted immediately after the stop codon in the 3ʹ UTR of *Nanog* (Li et al., 2019). In contrast, in this study, the RPB1 and BRD4 clusters were equally proximal to the mTetR spots in the ON state. This difference can be explained by the different insertion sites of the reporter sequences. In the STREAMING-tag system, MCP spots were anticipated to appear soon after the transcription of the *Nanog* start codon. Since the MS2 repeats are possibly co-transcriptionally spliced out through the SA and SD within the STREAMING-tag, the MCP spots were observed only when the 5ʹ-proximal region was transcribed. Therefore, most of the elongating RNAPII complexes may not be in the immediate vicinity of MS2-TetO repeats and mTetR in the STREAMING-tag system (Fig. 7D). As we analyzed the RF cluster nearest to mTetR, RPB1 clusters in the initiation complexes could be primarily detected rather than elongating RPB1 clusters in downstream regions. Thus, a detailed analysis of RF clusters along the gene length may be possible by the combined use of the STREAMING-tag and the standard MS2 systems to assess the transcriptional activity near the TSS and at the transcription termination site, respectively.

In addition, the tag can be visualized in both the ON and OFF states, while the gene position is detectable only in the ON state when the MS2 system is independently used. In fact, it has been demonstrated using the MS2 system and smFISH that RFs such as RPB1, BRD4, and Mediator form clusters in the nucleus and co-localize with transcriptionally active genes (Li et al., 2019, 2020; Sabari et al., 2018). However, the characteristics of the OFF state cannot be analyzed using these systems. Since the formation and dissolution of RF assemblies is anticipated to coordinate with the transcription activity (Henninger et al., 2021), identifying the gene and the associated RFs in the OFF state is crucial to understand the spatial relationship between specific genes and various RF clusters in different states.

### Detecting RNAPII Ser5ph in living cells using a mintbody

In this study, we developed a genetically encoded mintbody that recognizes RNAPII Ser5ph in living cells. As mintbodies repeatedly bind to and dissociate from their respective targets in living cells, they have no effect on cell division, embryo development, and differentiation (Sato et al., 2021). The formation of RNAPII Ser5ph-specific mintbody foci was sensitive to the CDK7 specific inhibitor THZ1 (Fig. S6G-J), suggesting that the presence of the mintbody does not block Ser5 dephosphorylation. The RNAPII Ser2ph-specific mintbody has recently been developed (Uchino et al., 2021). Thus, the specific RNAPII forms in the initiation and elongation complexes can now be visualized, which will facilitate future studies regarding transcriptional regulation in living cells.

### Spatial relationship between *Nanog* and RF clusters in the ON and OFF states

Using the STREAMING-tag system, we observed distinct association profiles of RF clusters and *Nanog* (mTetR) in the ON and OFF states (Fig. 7D). The ON state of *Nanog* is associated to the nearest RPB1 and BRD4 clusters with a proximity <350 nm, whereas the nearest MED19 and MED22 clusters are consistently proximal (<350 nm) to *Nanog*, regardless of their transcriptional states.

In addition to RPB1, its Ser5- and Ser2-phosphorylated forms were also localized in the vicinity of *Nanog* in the ON state (Fig. 7D). This observation is consistent with the view that transcription complexes containing RNAPII are assembled upon initiation of a transcriptional burst of *Nanog*, rather than significant pools of paused or poised RNAPII being associated with the gene (e.g., *Drosophila* heat-shock, mammalian *β-globin*, *c-myc*, and *c-fos*) during the OFF state (Gariglio et al., 1981; Gilmour and Lis, 1986; Plet et al., 1995). The RPB1 spots nearest to mTetR can represent either unphosphoylated, Ser5ph, or Ser2ph forms. However, since only the nearest RPB1 spot was analyzed, most spots were more likely to represent the unphosphorylated or Ser5ph forms than Ser2ph, whose foci are furthest from mTetR. The BRD4 clusters, which are often associated with enhancers were also closer to *Nanog* in the ON state. MED19 and MED22 clusters that are closely located near *Nanog*, independent of the transcriptional state, may serve as a scaffold for the formation of new initiation clusters containing BRD4 and hypophosphorylated RNAPII.

### Limitations

The STREAMING-tag is 5.5 kb long and contains essential components, such as MS2 repeat, TetO repeat, and an intein with G418 resistance protein. Hence, its insertion into the TSS-proximal region may affect gene expression. We observed that monoallelic knock-in of STREAMING-tag in *Nanog*, *Sox2*, and *Usp5* caused a slight decrease in protein expression (Fig. 2C, S2). Since protein splicing occurred quite efficiently in NSt cells, it is likely that the decrease in mRNA expression led to a concomitant decrease in protein expression levels. As intron size negatively correlates with the gene expression level (Castillo-Davis et al., 2002; Marais et al., 2005; Swinburne and Silver, 2008), the insertion of a STREAMING-tag containing one intron could slow down the overall RNA production via transcription elongation and splicing. However, remarkably, there was no significant difference in the percentage of cells with visible transcriptional spots between knock-in and non-knock-in alleles. This suggests that although the knock-in of the STREAMING-tag moderately decreased gene expression, it did not significantly affect the initiation, pause-release, and productive elongation events. Thus, the STREAMING-tag system can be a useful technique for analyzing promoter proximal transcription as a result of initiation and pause-release. Therefore, we recommend a monoallelic knock-in because the optimum expression levels may be critical for the function of certain genes.

Since the STREAMING-tag system requires knock-ins, the technical problems associated with the knock-in system also apply to the STREAMING-tag. For example, knock-in involving non-expressed genes requires a selectable marker with a constitutive promoter and an excision system, such as the Cre-loxP system, and obtaining the desired cell line is effort-intensive. Recently, it has been reported that dCas13 and dCas9 can be used to simultaneously visualize the transcription and genomic loci of target genes without knock-in (Wang et al., 2019). However, this method requires the introduction of fluorescently labeled proteins or RNAs directly into cells, which is not suitable for long-term live cell imaging. In addition, labeling non-repeat sequences with dCas9 or dCas13 requires many types of sgRNAs (Gu et al., 2018; Li et al., 2020).

In addition, in the STREAMING-tag, we used the TetO/TetR system to label DNA regions making it impossible to combine with the tet-inducible expression system (such as Tet-ON and Tet-OFF) (Das et al., 2016). To circumvent this issue, the LacO/LacI and ParS/ParB systems may be used as alternatives (Saad et al., 2014; Tsukamoto et al., 2000).

## Acknowledgments

We thank Ms. Yuki Ochiai for providing technical assistance. This work was supported by Grants-in-Aid for Scientific Research from the Ministry of Education, Culture, Sports, Science, and Technology (JP18H05531, JP19K06612, and JP21H05753 to H.Oc., JP21H04764 to H.K., JP20K06484 to Y.S, and JP18H05527 to Y.O. and H.K.) and the JST CREST program (JPMJCR16G1 to Y.O., H.K., and H.Oc.; and JPMJCR20S6 to Y.S).

## Author contributions

H.Oc. conceived, designed, and supervised the study and established all the knock-in cells. H.Oc. and S.S. performed verification experiments on the basic design of the STREAMING-tag. H.Oc., H.Oh., S.S., M.S., and H.Ow. established mintbody-SNAPtag and NLS-SNAPtag expressing cells and performed the MSCD experiments. H.Oh. performed STREAMING-tag and SNAPtag imaging experiments and analyzed the data. J.L. and A.P. developed the *Rpb1-*, *Brd4-*, *Med19-*, and *Med22*-SNAPtag knock-in vectors and their corresponding CRISPR vectors. S.U., Y.S., Y.O., and H.K. developed the RNAPII Ser5ph mintbody and performed validation experiments. H.K. supervised the development of mintbodies. H.Oc, H.Oh, T.Y., A.P., and H.K. interpreted the data and wrote the manuscript.

## STAR methods

### KEY RESOURCES TABLE

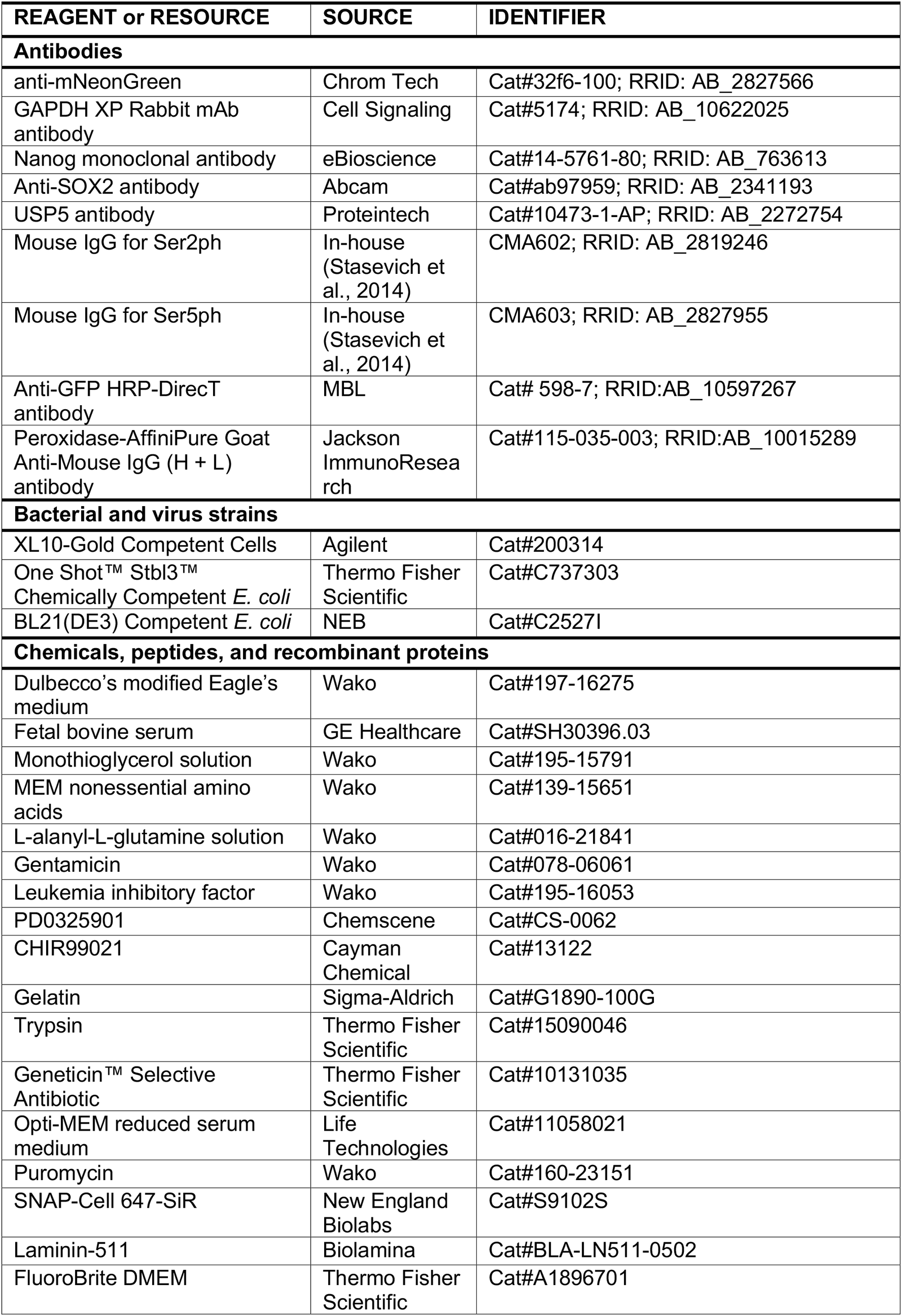
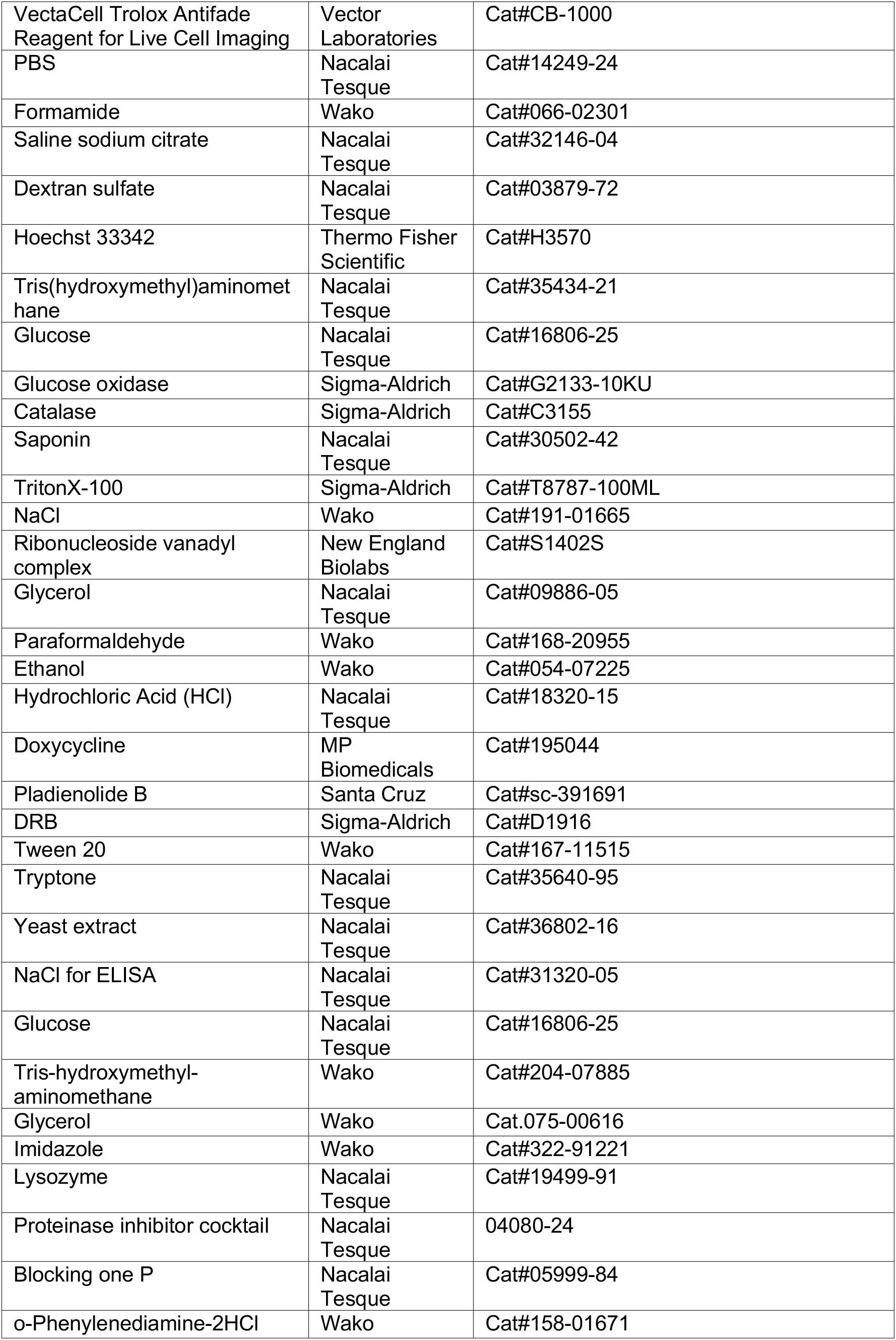
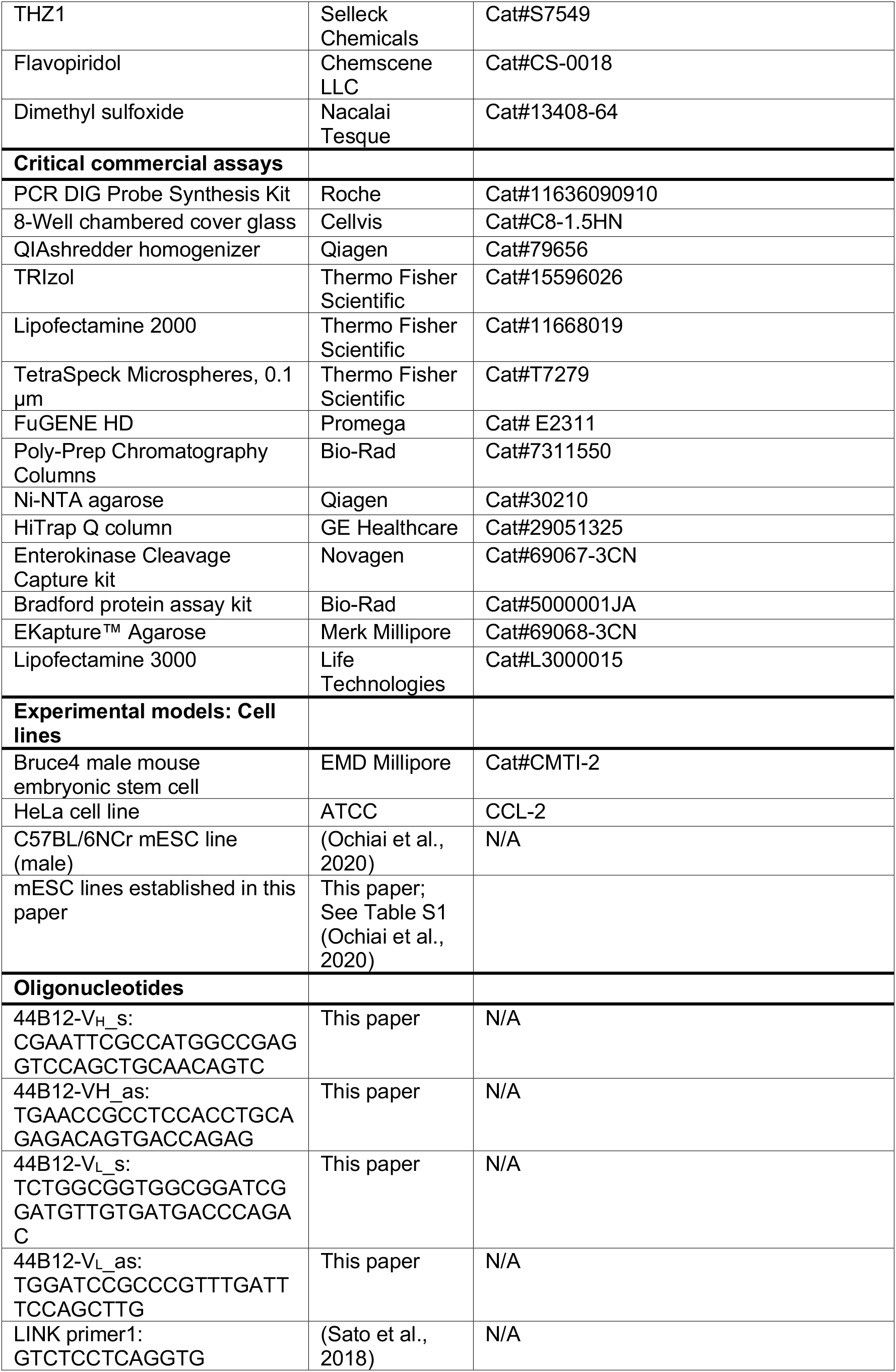
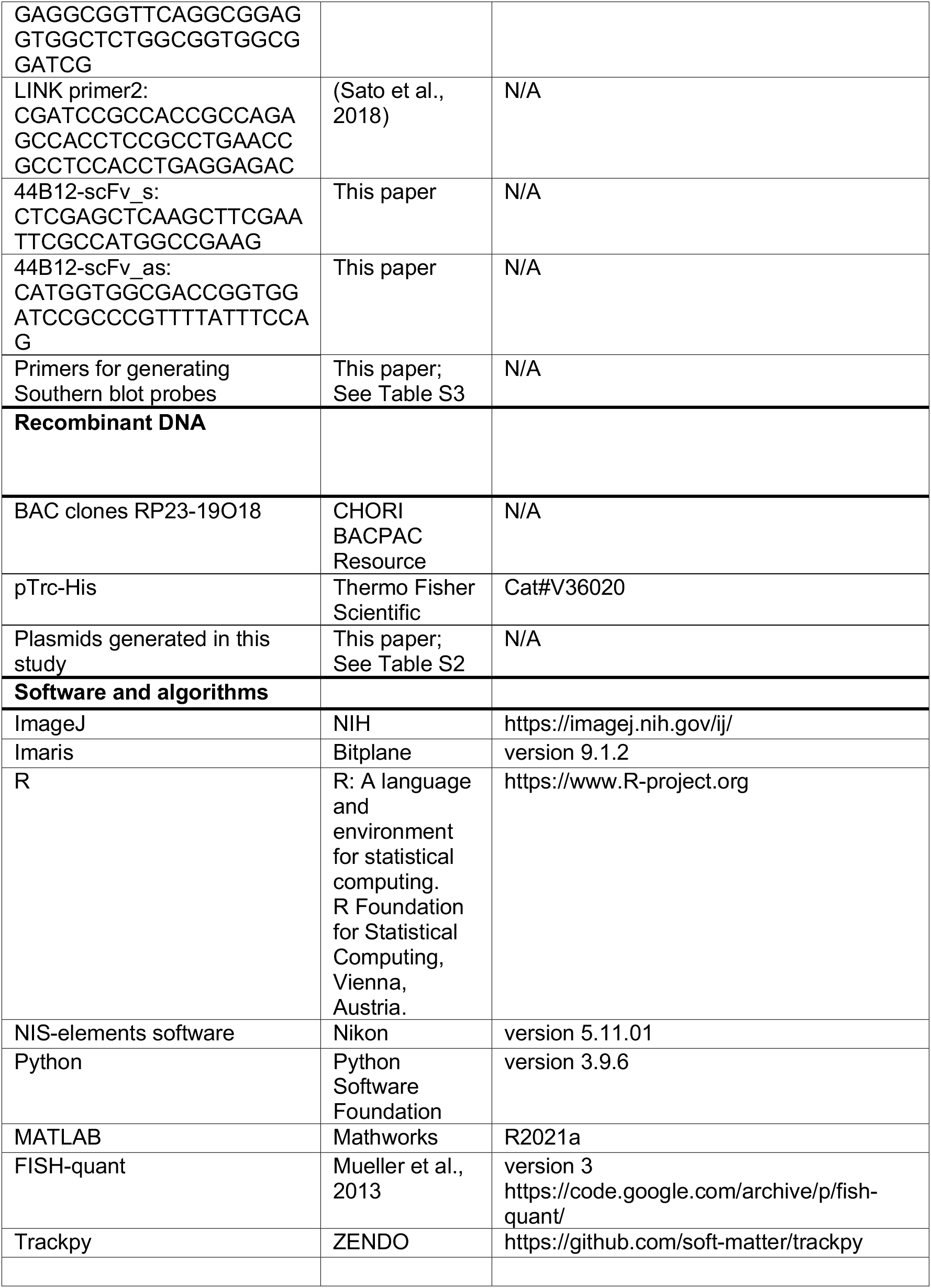

## RESOURCE AVAILABILITY

### Lead contact

Further information and requests for resources and reagents should be directed to and will be fulfilled by the lead contact Hiroshi Ochiai.

### Materials availability

Plasmids generated in this study have been deposited to Addgene (https://www.addgene.org/Hiroshi_Ochiai/).

### Data and code availability

This study does not report the original code. Any additional information required to reanalyze the data reported in this paper is available from the lead contact upon request.

## EXPERIMENTAL MODEL AND SUBJECT DETAILS

### Cell lines

Wild-type (WT) mESCs derived from inbred mice (Bruce 4 C57BL/6J, male, EMD Millipore, Billerica, MA, USA) and other knock-in derivatives were cultured as described previously (Ochiai et al., 2014). C57BL/6NCr (male) mESCs (Ochiai et al., 2020) were used as *Sox2* STREAMING-tag knock-in cells. Briefly, all mESC lines were maintained in 2i medium (Dulbecco’s modified Eagle’s medium [DMEM, Wako, Osaka, Japan, 197-16275]; 15% fetal bovine serum [FBS, GE Healthcare, Little Chalfont, UK, SH30396.03]); 0.5 mM monothioglycerol solution [Wako, 195-15791]; 1×MEM nonessential amino acids [Wako, 139-15651]; 2 mM L-alanyl-L-glutamine solution [Wako, 016-21841]; 1,000 U/mL leukemia inhibitory factor [Wako, 195-16053]; 20 µg/mL gentamicin [Wako, 078-06061]; 3 µM CHIR99021 [Cayman Chemical, Ann Arbor, MI, USA, 13122]; and 1 µM PD0325901 [Chemscene, Monmouth Junction, NJ, USA, CS-0062]) on a 0.1% gelatin (Sigma-Aldrich, St. Louis, MO, USA, G1890-100G)-coated dish at 37°C and 5% CO_2_. The cell lines used in this study are listed in Table S1.

HeLa cells were grown in DMEM high-glucose medium (Nacalai Tesque, Kyoto, Japan) containing 10% FBS (Gibco, Grand Island, NY, USA) and 1% L-glutamine – penicillin– streptomycin solution (GPS; Sigma-Aldrich, St. Louis, MO, USA) at 37°C in a 5% CO_2_ atmosphere.

## METHOD DETAILS

### Plasmids

Plasmids were constructed using common molecular biological techniques. A list of plasmids used can be found in Table S2 and is available from Addgene (Watertown, MA, USA; https://www.addgene.org/Hiroshi_Ochiai/).

### Optimization of selection marker flanked by split inteins

C57BL/6J mESC cell lines (2.5 × 10^5^) were plated into each well of a 24-well plate and transfected with 50 ng pCAG-hyPBase (Ochiai et al., 2015) and 450 ng Intein-related plasmids (Tables S1 and S2) using Lipofectamine 3000 (L3000015, Life Technologies, Carlsbad, CA, USA) according to the manufacturer’s instructions. After 24 h, the medium was replaced with fresh medium. After another 24 h, the cells were detached using 0.25% trypsin (Thermo Fisher Scientific, Waltham, MA, USA, 15090046) with 1 mM EDTA and plated into two wells of a 24-well plate, each containing 1.25 × 10^5^ cells; one of the wells was treated with 200 µg/mL G418 (Geneticin™ Selective Antibiotic, Thermo Fisher Scientific, 10131035). After 2 days, the medium was replaced with fresh medium containing 300 µg/mL G418. After 24 h, the number of surviving cells was counted. Cells expressing PInt-I and CInt-I were passaged and seeded into an 8-well chambered cover glass with #1.5 glass (Cellvis, Sunnyvale, CA, USA, C8-1.5HN). On the next day, images were acquired using a Nikon Ti-2 microscope (Nikon, Tokyo, Japan) with a CSU-W1 confocal unit (Yokogawa Electric, Tokyo, Japan), a 20× Nikon Plan Fluor objective lens (NA 0.5), and an iXon Ultra EMCCD camera (Andor Technology, Belfast, UK, DU-888U3-CSO-#BV), with laser illumination at 488 nm, and were analyzed using NIS-elements software (version 5.11.01, Nikon); 41 z planes per site spanning 20 µm were acquired. These cells were further expanded, and proteins were extracted from the cells for western blot analysis.

### STREAMING-tag knock-in by genome editing

First, we describe the establishment of *Nanog* STREAMING-tag knock-in cells. C57BL/6J mESCs (5 × 10^5^) were plated into each well of a 12-well plate; after 1 h, the following transfection reagents were mixed: 2 µg of targeting vector (e.g., pTV-Nanog_2-5prime-1000-24MS96T-3F_NeoR), 700 ng of CRISPR vector (e.g., eSpCas9-EF-5Nanog_2), and 300 ng of pKLV-PGKpuro2ABFP. Subsequently, 62.5 µL of Opti-MEM-reduced serum medium (Life Technologies, 11058021) and 2.5 µL of P3000 reagent (Life Technologies, L3000015) were added to each plate. In a separate tube, 62.5 µL of Opti-MEM-reduced serum medium and 4.5 µL of Lipofectamine 3000 (Life Technologies, L3000015) were added per reaction and mixed well. The P3000 and Lipofectamine 3000 media were mixed in equal volumes and incubated for 15 min at room temperature (RT). The complex was then added to the wells containing the cells and incubated overnight. After 24 h, the medium was replaced with fresh 2i medium containing 1 µg/mL puromycin (Wako, 160-23151) to eliminate untransfected cells. After 24 h, the medium was replaced with fresh 2i medium; 24 h later, all cells were transferred to gelatin-coated 10 cm dishes. After 48 h, the medium was replaced with 2i medium containing 200 µg/mL G418; 48 h later, the medium was replaced with 2i medium containing 200 µg/mL G418. After another 48 h, 24 colonies were selected for further analysis. Genomic DNA was extracted from these cells, and genomic PCR was performed to narrow down the candidate cell lines. Thereafter, candidate clones were analyzed using Southern blotting, as described previously (Ochiai et al., 2014). The restriction enzymes and genomic regions used for the Southern blot probes are summarized in Table S3. Probes were prepared using the PCR DIG Probe Synthesis Kit (Roche, Basel, Switzerland, 11636090910).

The same procedure was used to knock-in the STREAMING-tag in *Pou5f1*, *Sox2*, *Wnk1*, *Flnc*, and *Usp5*. The plasmids used are listed in Tables S1 and S2, respectively.

### Establishment of fluorescent protein-expressing cells using the piggyBac system

NSt-GR cells were established as follows: NSt mESCs (2.5 × 10^5^) were plated into each well of a 24-well plate, and after 1 h, the following transfection reagents were mixed. In a tube, 50 ng pCAG hyPBase and 75 ng mTetR-mNG expression vectors (such as pLR5-CAG-TetR_W43F-3xmNG) and 375 ng of pLR5-CAG-hMCP-mScarlet-I-NLS were mixed. Next, 25 µL of reduced serum Opti-MEM and 1 µL of P3000 reagent were added to each plate. In a separate tube, 25 µL of reduced serum Opti-MEM and 1.8 µL of Lipofectamine 3000 were added per reaction and mixed well. The P3000 and Lipofectamine 3000 media were mixed in equal volumes and incubated at RT for 15 min. The medium was replaced with fresh 2i medium after 24 h (day 2).

Every 24 h, the medium was replaced with fresh 2i medium. On day 5, all cells were passaged in 12-well plates, and the cells were collected on day 6 following treatment with trypsin. Cells moderately expressing mTetR-mNG and MCP-RFP were isolated using a BD FACSAria III cell sorter (BD Biosciences, Franklin Lakes, NJ, USA) and plated into gelatin-coated 6 cm dishes (Supplementary Methods 1, Fig. S7). At 8 days after fluorescence-activated cell sorting (FACS), colonies were picked and plated into a gelatin-coated 8-well chambered cover glass. Three days later, cells expressing moderate amounts of fluorescent protein, referred to as NSt-GR cells, were observed using fluorescence microscopy and used for further experiments.

For transfection with transgene containing SNAPtag, the following method was used: NSt mESC or other cell lines (2.5 × 10^5^) were plated into each well of a 24-well plate, and after 1 h, the following transfection reagents were mixed: 50 ng pCAG hyPBase, 75 ng pLR5-CAG-TetR_W43F-3xmNG, 275 ng pLR5-CAG-hMCP-mScarlet-I-NLS, and 100 ng of SNAPtag expression vector (e.g., pLR5-CAG-NLS-SNAP). To each of these, 25 µL of reduced serum Opti-MEM and 1 µL of P3000 reagent were added. In a separate tube, 25 µL of Opti-MEM reduced serum medium and 1.8 µL of Lipofectamine 3000 were added per reaction and mixed well. The P3000 and Lipofectamine 3000 media were mixed in equal volumes and incubated at RT for 15 min. After 24 h (day 2), the medium was replaced with 2i medium. After another 24 h (day 3), the medium was replaced with 2i medium, and the cells were passaged in 12-well plates on day 5. On day 6, the cells were incubated in 2i medium containing 300 nM SNAP-Cell 647-SiR (New England Biolabs, Ipswich, MA, USA, S9102S) for 30 min at 37°C and 5% CO_2_. The cells were washed three times with 2i medium and incubated at 37°C and 5% CO_2_ for another 30 min. The cells were collected following treatment with trypsin, and cells moderately expressing mTetR-mNG, MCP-RFP, and SNAPtag were sorted using a BD FACSAria III cell sorter and seeded into gelatin-coated 6 cm dishes (Supplementary Methods 1, Fig. S7). The medium was changed once every 2 days. At 8 days after FACS sorting, colonies were picked and plated into a gelatin-coated 8-well chambered cover glass and cultured. After 3 days, cells expressing a moderate amount of fluorescent protein were observed under a fluorescence microscope and used for further experiments. The cells were expected to show a mild level of fluorescence expression; if the expression level was too high, it was difficult to detect spots and foci (Supplementary Methods 1, Fig. S7).

### RF-SNAPtag knock-in

NSt-GR mESCs (1.25 × 10^5^) were plated into each well of a 24-well plate, and after 1 h, the transfection reagents were mixed. In a tube, 500 ng of targeting vector (e.g., Rpb1 snap targeting vector), 250 ng of CRISPR vector (e.g., eSpCas9-Rpb1-gRNA), and 75 ng of pKLV-PGKpuro2ABFP were mixed. To each of these, 31 µL of reduced serum Opti-MEM and 1.25 µL of P3000 reagent were added. In a separate tube, 31 µL of reduced serum Opti-MEM and 2.25 µL of Lipofectamine 3000 were added per reaction and mixed well. The P3000 and Lipofectamine 3000 media were mixed in equal volumes and incubated at RT for 15 min. After 24 h (day 2), the cells were treated with 1 µg/mL puromycin in 2i medium. The medium was replaced with fresh 2i medium after another 24 h (day 3). Every 24 h, the medium was replaced with fresh 2i medium. On day 6, the cells were incubated for 30 min in 2i medium containing 300 nM SNAP-Cell 647-SiR at 37°C and 5% CO_2_. The cells were washed three times with 2i medium and incubated at 37°C and 5% CO_2_ for another 30 min. The cells were collected by trypsin treatment, and SNAPtag signal-positive cells were sorted using a BD FACSAria III cell sorter and seeded into gelatin-coated 6 cm dishes. The medium was changed once every 2 days. Twenty-four colonies were picked on day 8 after FACS. Genomic DNA extracted from these cells was used for genomic PCR to narrow down the candidate cell lines. Candidate clones were further analyzed using Southern blotting, as described previously (Ochiai et al., 2014). The restriction enzymes and genomic regions used for the Southern blot probes are summarized in Table S3. Probes were prepared using a PCR DIG Probe Synthesis Kit (Roche Diagnostics, Mannheim, Germany).

The SNAPtag was knocked into *Brd4*, *Med19*, and *Med22*, as described above. See Tables S1 and S2 for the plasmids used in this study.

### Microscopy

For live cell imaging of mESCs, the medium was replaced with imaging 2i medium (FluoroBrite DMEM [Thermo Fisher Scientific, A1896701], 15% FBS [GE Healthcare, SH30396.03], 0.5 mM monothioglycerol solution [Wako, 195-15791], 1× MEM nonessential amino acids [Wako, 139-15651], 2 mM L-alanyl-L-glutamine solution [Wako, 016-21841], 1,000 U/mL LIF [Wako, 195-16053], 20 µg/mL gentamicin [Wako, 078-06061], 3 µM CHIR99021 [Cayman Chemical, 13122], 1 µM PD0325901 [Chemscene, CS-0062], and VectaCell Trolox Antifade Reagent for Live Cell Imaging [Vector Laboratories, Burlingame, CA, USA, CB-1000; 1:1,000]). For single-molecule fluorescence *in situ* hybridization (smFISH), the samples were mounted in catalase/glucose oxidase-containing mounting media (GLOX; 0.4% glucose [Nacalai Tesque, 16806-25] in 10 mM Tris-HCl [pH 8.0], 2× saline sodium citrate [SSC], glucose oxidase [37 µg/mL, Sigma-Aldrich, G2133-10KU], and 1/100 catalase [Sigma-Aldrich, C3155]). Images were acquired using a Nikon Ti-2 microscope with a CSU-W1 confocal unit, a 100× Nikon Apo TIRF oil-immersion objective lens (NA 1.49), and an iXon Ultra EMCCD (Andor Technology), operated using NIS-Elements software (ver. 5.11.01; Nikon). The microscope was also equipped with 405, 488, 561, and 637 nm lasers (Andor Technology), a stage-top microscope incubator for live cells (5% CO_2_; 37°C; STXG-TIZWX-SET, Tokai Hit, Shizuoka, Japan), and an ASI MS-2000 piezo stage (ASI). Z-stack images spanning 20 µm with 200 nm intervals (101 sections; 130 nm/pixel) were acquired.

### Snapshot fluorescence imaging of live cells

Cells (5 × 10^4^) were plated onto each well of an 8-well chambered cover glass (Cellvis) that was pre-coated with laminin-511 (BioLamina, Sundbyberg, Sweden, BLA-LN511-0502) and cultured overnight. For imaging SNAPtag, the cells were incubated in 2i medium containing 300 nM SNAP-Cell 647-SiR for 30 min at 37°C and 5% CO_2_, washed three times with fresh 2i medium, and incubated for another 30 min at 37°C and 5% CO_2_. The medium was then replaced with imaging 2i medium (FluoroBrite DMEM [Thermo Fisher Scientific, A1896701] containing VectaCell Trolox Antifade Reagent for Live Cell Imaging (Vector Laboratories, CB-1000; 1:1,000). After image acquisition, the images were processed with a one-pixel diameter 3D median filter using ImageJ software (NIH, Bethesda, MD, USA).

### Western blotting

Cells were washed twice with phosphate-buffered saline (PBS, Nacalai Tesque, 14249-24), trypsinized, and collected by centrifugation at 190 × g for 2 min at 20°C . The cells were counted and washed twice with PBS. The cells were then lysed in lysis buffer (0.5% Triton X-100 (Sigma-Aldrich, T8787-100ML), 150 mM NaCl (Wako, 191-01665), and 20 mM Tris-HCl [pH 7.5]) to obtain 2 × 10^6^ cells per 100 µL. The lysates were then incubated at 95°C for 5 min and filtered using a QIAshredder homogenizer (Qiagen, 79656). The extracted proteins were analyzed using 5–20% gradient sodium dodecyl sulfate-polyacrylamide gel (SDS-PAGE) electrophoresis and transferred onto Immobilon Transfer Membranes (Millipore, INYC00010) for immunoblotting. The primary antibodies used were anti-mNeonGreen (1:500; Chrom Tech, Apple Valley, MN, USA, 32f6-100, RRID:AB_2827566), anti-GAPDH (1:5000; Cell Signaling Technology, Danvers, MA, USA, 5174, RRID:AB_10622025), anti-NANOG (1:1000; eBioscience, San Diego, CA, USA, 14-5761-80, RRID:AB_763613), anti-SOX2 (1:1000; Abcam, Cambridge, UK, ab97959, RRID:AB_2341193), and anti-USP5 (1:2000; 10473-1-AP, Proteintech, Rosemont, IL, USA, RRID:AB_2272754).

### smFISH

Cells (5 × 10^4^) were transferred onto a laminin 511-coated 8-well chambered cover glass and cultured for 1 h at 37°C and 5% CO_2_. The cells were washed with PBS, fixed with 4% paraformaldehyde (Wako, 168-20955) in PBS for 10 min, and washed twice with PBS. The cells were then permeabilized in 70% ethanol (Wako, 054-07225) at 4°C overnight. After washing with 10% formamide (Wako, 066-02301) dissolved in 2× SSC (Nacalai Tesque, 32146-04) buffer, the cells were hybridized to probe sets in 130 µL of hybridization buffer containing 2× SSC, 10% dextran sulfate (Nacalai Tesque, 03879-72), 10% formamide, and 1 µM TetR probe (5′-TCCCTATCAGTGATAGAGANTATTCGGCTCCGCGCGCGGAGCCGAATACCT CGC-3′). Hybridization was performed for 12 h at 37°C in a moist chamber. The coverslips were washed with 10% formamide in 2× SSC solution and incubated at 37°C for 30 min in the dark. The cells were hybridized to probe sets in 130 µL of hybridization buffer containing 2× SSC, 10% dextran sulfate, 10% formamide, and 125 nM Alexa 488 probe (5′-TGCGAGGTATTCGGCTCCGCGT-3′). Both ends were modified with Alexa 488 and *Nanog* Exonic probes (Ochiai et al., 2014). Hybridization was performed for 4 h at 37°C in a moist chamber. The coverslips were washed with 10% formamide in 2× SSC solution, incubated at 37°C for 30 min in the dark, and then washed with 10% formamide in 2× SSC solution with Hoechst 33342 (1:1000, Thermo Fisher Scientific, H3570) and incubated at 37°C for 30 min in the dark. Hybridized cells were mounted in GLOX buffer. After image acquisition, the images were filtered with a one-pixel-diameter three-dimensional median filter and subjected to background subtraction via a rolling ball radius of 5 pixels using ImageJ software. Detection and counting of smFISH signals were performed using MATLAB FISH-quant software version 3 (Mueller et al., 2013). For transcriptional spot detection, the processed images were subjected to maximum-intensity projection. Transcriptional spots were detected using the projected images and ComDet plug-in of ImageJ (https://github.com/ekatrukha/ComDet/wiki).

### FRAP analysis

NSt-MT(WT) and NSt-MT(W43F) cells (Table S1) (5 × 10^4^) were plated into each well of laminin-511-coated 8-well chambered cover glass and cultured overnight. The medium was replaced with an imaging medium (2i). For photobleaching, a fluorescence recovery after photobleaching (FRAP) module (Nikon) was used in combination with a CSU-W1 confocal system. Five z-stack images (17 sections at 0.2 µm intervals) were taken at 4 s intervals, and after applying a 488 nm laser pulse (100 ms; 24.2% laser attenuation; 50 mW laser output) through a FRAP module, 25 z-stack images (17 sections at 0.2 µm intervals) were taken at 4 s intervals. The images were processed with a one-pixel-diameter 3D mean filter and subjected to maximum intensity projection and “bleach correction” using ImageJ software. After manually selecting the center of the TetR regions, the average fluorescence intensities of a circular region of interest of 6 pixels in diameter with the center were measured.

The normalized intensity *I*_norm_ was calculated using the following equation:

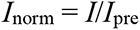

where, *I_pre_* is the fluorescence intensity of pre-FRAP. To extract the characteristic timescales of fluorescence recovery and the mobile fractions, average FRAP curves were fitted using R with the following function:

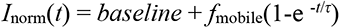

where, *baseline*, *τ*, and *f*_mobile_ represent the expected *I*_norm_(0), recovery time constant, and mobile fraction, respectively. Assuming that the *baseline* was the same for TetR(WT) and TetR(W43F)/mTetR, the sum of the squares of the offsets (*S*) was used to estimate the best-fit curve. *S*WT + *S*W43T was minimized when the *baseline* value was 0.409.

## DNA-FISH

The cells were plated onto laminin-511-coated glass slides and cultured overnight at 37°C and 5% CO_2_. The cells were washed with PBS, fixed with 4% paraformaldehyde in PBS for 10 min, washed twice with PBS, and then treated with PBS containing Hoechst 33342 nucleic acid stain (1:1000) for 10 min. Images were acquired using a Nikon Ti-2 microscope with a CSU-W1 confocal unit, a 100× Nikon Apo TIRF oil-immersion objective lens (NA 1.49), and an iXon Ultra EMCCD camera with laser illumination at 405, 488, and 637 nm. In this setup, the pixel size was 130 nm, and 76 z-planes per site spanning 15 µm (z-step = 200 nm) were acquired. The cells were then subjected to 3D-DNA-FISH as previously described with some modifications (Bolland et al., 2013). Briefly, the cells were washed twice with PBS and then permeabilized in 0.1% saponin (Nacalai Tesque, 30502-42)/0.1% Triton X-100/2 mM ribonucleoside vanadyl complex (New England Biolabs, S1402S) in PBS for 10 min at RT. Following two washes with PBS, the cells were incubated for 20 min in 20% glycerol/PBS at RT and stored in 50% glycerol/PBS at −20°C for at least 1 day. After incubation, the cells were recalibrated at RT in 20% glycerol/PBS and subjected to three successive freeze/thaw cycles in liquid nitrogen (Bolland et al., 2013). Thereafter, the cells were washed twice with PBS for 5 min each at RT, incubated in 0.1 M HCl (Nacalai Tesque, 18320-15) for 30 min at RT, washed once again with PBS for 5 min at RT, permeabilized in 0.5% saponin/0.5% Triton X-100 in PBS for 30 min at RT, washed two more times with PBS for 5 min per wash at RT, and then equilibrated in 50% formamide/2× SSC for 10 min at RT. Next, the cells were hybridized to a pre-denatured *Nanog* probe (see below) using a hybridization buffer containing 1× SSC, 10% dextran sulfate, and 50% formamide. Hybridization was performed for 16 h at 37°C in a moist chamber. The cells were washed in 2× SSC for 5 min at RT, 50% formamide/2× SSC for 15 min at 45°C, 2× SSC for 5 min at 45°C, and then 2× SSC for 5 min at RT, followed by a wash in 2× SSC with Hoechst 33342 (1:1000) for 10 min at RT. Hybridized cells were mounted in GLOX buffer (Ochiai et al., 2014). Images were acquired using a laser illumination set at 405 nm for Hoechst 33342 and at 647 nm for *Nanog* probes. The BAC clones RP23-19O18 (CHORI BACPAC Resources, Emeryville, CA, USA) were used as DNA-FISH probes for *Nanog*.

### SNR analysis

The NMP-R mESCs (Ochiai et al., 2015) (5 × 10^4^) were plated into 24-well plates on the day before transfection. The cells were transfected with 700 ng of MS2 sgRNA expression vectors (Ochiai et al., 2015) on the following day using Lipofectamine 2000 (Thermo Fisher Scientific, 11668019). After 12 h, the cells were treated with puromycin (2 µg/mL) and doxycycline (100 ng/mL, MP Biomedicals, Santa Ana, CA, USA, 195044) for another 24 h. The cells were trypsinized, transferred onto a laminin-511-coated 8-well chambered cover glass (CellVis, C8-1.5H-N), and cultured overnight at 37°C and 5% CO_2_ in doxycycline-containing medium.

For NSt-GR cells, an 8-well chambered cover glass (CellVis, C8-1.5H-N) coated with laminin-511 was used. Cells (5 × 10^4^) were plated into each well and cultured at 37°C and 5% CO_2_ overnight.

After image acquisition, the area outside the cell nucleus was measured as the average background intensity, and the intensity was subtracted from the entire image using ImageJ software. The standard deviation (*σN*) was measured in the nucleus. Next, the DNA-labeled region was selected using the “Find Maxima” function of TrackMate with an estimated blob diameter of 0.5 µm, and the mean intensity value (*µ*) of the target foci was measured. The signal-to-noise ratio (SNR) was calculated using the following equation:

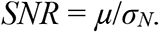

### MSCD analysis

NSt-NLS-SNAP and PSt-NLS-SNAP mESCs (5 × 10^4^) were plated into an 8-well chambered cover glass with laminin-511 and cultured overnight at 37°C and 5% CO_2_. The cells were incubated in 2i medium containing 300 nM SNAP-Cell 647-SiR for 30 min at 37°C and 5% CO_2_. The cells were then washed three times with 2i medium and incubated for another 30 min at 37°C and 5% CO_2_. After the medium was replaced with imaging 2i medium, 46 z-sections spanning 9 µm (z-step = 200 nm) were acquired at 15 s intervals for 450 s. The acquired images were filtered with a one-pixel diameter 3D Gaussian Blur filter using ImageJ software. The mTetR-mNG images were subjected to background subtraction using a rolling ball radius of five pixels using ImageJ software. Fluorescent spots were detected using “Spot” function of Imaris software (version 9.1.2, Bitplane, Zürich, Switzerland) with the spot diameter set to 0.8 µm (semi-automatic detection). The nucleus center of mass was determined from NLS-SNAP fluorescence using ImarisCell (Bitplane), with the Cell Smooth Filter Width and Cell Background Subtraction Width parameters set at 1 and 0.64 µm, respectively. The mean square change in distance (MSCD) was calculated as the average change in distance between the nuclear center of mass and genomic locus over all possible combinations of time points separated by the lag time *Δt*; *MSCD* = [*d*(*t*)−*d*(*t*+*Δt*)]^2^ (Arai et al., 2017; Meister et al., 2010). To categorize the “ON” and “OFF” states, the threshold was set based on the histogram of fluorescence intensities in the mTetR-mNG area in the MCP channel of all analyzed images. A valley was typically found between a peak at the background level and broad peaks at higher intensities, and the value at the valley (*fluorescence intensity* = 5) was used as the threshold. If transcription was observed at least once during the time-lapse, the data were classified as “ON.”

### DRB release assay

NM-G (Ochiai et al., 2014) and NSt-NLS-SNAP cells (9 × 10^4^) were seeded onto a laminin 511-coated 8-well chambered cover glass and cultured overnight. The medium was replaced with 2i medium containing 100 µM 5,6-dichloro-1-β-d-ribofuranosylbenzimidazole (DRB, Sigma-Aldrich, D1916) and incubated for 1.5 h. The cells were then washed three times with fresh 2i medium. The medium was replaced with an imaging medium (2i), and 51 z-sections spanning 20 µm (z-step = 200 nm) were acquired at 1 min intervals for 30 min. The images were processed using ImageJ with Gaussian Blur 3D (pixel size = 1), background subtraction with a rolling ball radius of 50 pixels, and maximum intensity projection. MS2 coat protein (MCP) spots were defined manually.

### Cloning antibody variable fragments encoding 44B12

Mouse hybridoma cells expressing Ser5ph-specific antibodies were generated (MAB Institute, Inc., Nagano, Japan) as previously described (Kimura et al., 2008), using a peptide Ser-Tyr-Ser-Pro-Thr-phosphoSer-Pro-phosphoSer-Tyr-Ser-Pro-Thr-Ser-Pro-Ser-Tyr-Ser-Pro-Cys, harboring Ser5ph and Ser7ph at the C-terminal domain sequence. The resulting antibody 44B12 reacted with peptides containing Ser5ph, regardless of the phosphorylation state of Ser7 (Fig. S6). To construct the mintbody, RNA was purified using TRIzol (Thermo Fisher Scientific, 15596026), and the sequences encoding IgG heavy and light chains were determined using RNA sequencing (Kuniyoshi et al., 2016). The variable regions of heavy and light chains (VH and VL) were each amplified using PCR with specific primers (KEY RESOURCES TABLE) and then connected using linker primers (KEY RESOURCES TABLE) as described previously (Sato et al., 2018; Tjalsma et al., 2021). The scFv fragment was cloned into the sfGFP-N1 vector (Addgene #54737) (Pédelacq et al., 2006) using In-fusion (Takara, Shiga, Japan) with the primers 44B12-scFv_s and 44B12-scFv_as; KEY RESOURCES TABLE) to generate a 44B12-sfGFP expression vector.

Two-point mutations predicted to improve the folding and/or stability of scFv were introduced into the RNAPII Ser5ph-mintbody as described previously (Uchino et al., 2021). Mintbodies were expressed in HeLa cells by transfecting the expression vectors using Fugene HD (Promega, Madison, WI, USA, E2311). A plasmid vector (2 µg) and Fugene HD (6 µL) were mixed in Opti-MEM (Thermo Fisher Scientific; 100 µL). After incubation at RT for 30 min, the mixture was added to HeLa cells grown in 35 mm glass-bottom dishes (AGC Technology Solutions, Kawasaki, Japan). The culture medium was changed to FluoroBrite DMEM containing 10% FBS and 1% GPS, and fluorescence images were acquired using a point-scan confocal microscope (Olympus, Tokyo, Japan, FV1000 with IX-81) with a UPlanApoN 60× OSC oil immersion objective lens (NA 1.4) using the built-in software FLUOVIEW ver. 4.2 (512 × 512 pixels, pixel dwell time 4.0 μs, pinhole 100 μm, zoom ×5.0, line averaging ×4, and a multi-argon ion 488 nm laser line with 10% transmission).

### Purification of RNAPII Ser5ph-mintbody

RNAPII Ser5ph-mintbody was purified using the same method as used for the Ser2ph-mintbody (Uchino et al., 2021). Briefly, a His-tag-RNAPII Ser5ph-mintbody expression vector was constructed using pTrc-His (Thermo Fisher Scientific, V36020). *Escherichia coli* BL21 (DE3) cells harboring the expression vector were grown in YTG medium (1% tryptone, 0.5% yeast extract, 0.5% NaCl, and 0.2% glucose; 100 mL) for 20 h at 18°C. After dilution in YTG medium (2 L), the cells were incubated for 8 h at 15°C, followed by incubation with isopropyl-β-D-thiogalactopyranoside (at a final concentration of 1 mM) for 12 h at 15°C. The cells were collected by centrifugation (4,000 ×*g*; 10 min; 4°C), and the pellet was stored at −80°C until thawing in 20 mL Buffer L (50 mM Tris-HCl, pH 8.0, 300 mM NaCl, 5% glycerol) containing 1 mg/mL lysozyme (Nacalai Tesque) and 1% proteinase inhibitor cocktail (Nacalai Tesque). After sonication (Branson Ultrasonics, Brookfield, CT, USA; Sonifier 250) for lysis, cell debris was removed by centrifugation (15,000 ×*g* for 15 min at 4°C). Ni-NTA agarose (Qiagen, Hilden, Germany; 0.5 mL) was equilibrated with Buffer L and settled in an open column (Poly-Prep Chromatography Columns; Bio-Rad, Hercules, CA, USA; 20 mL) before the cell lysate was applied at 4°C. After washing the column twice with Buffer L (10 mL each), elution buffer (Buffer L containing 150 mM imidazole, pH 8.0) was added (1 mL, three times). The eluted fractions were dialyzed against starting buffer (10 mM Tris-HCl, pH 7.0, 50 mM NaCl; 1 L) overnight with buffer exchange. The His-RNAPII Ser5ph mintbody was further purified using a HiTrap Q column (GE Healthcare) with a linear gradient elution with End Buffer (10 mM Tris-HCl, 1 M NaCl) using AKTAprime plus (GE Healthcare) at 4°C. After 10–20% SDS-PAGE and Coomassie Blue staining, fractions containing His-RNAPII Ser5ph-mintbody were pooled. After the His-tag was removed using an enterokinase cleavage capture kit (Novagen, Madison, WI, USA), His-tag- and enterokinase-free RNAPII Ser5ph-mintbody was prepared by passing through EKapture™ Agarose Millipore).

## ELISA

Enzyme-linked immunosorbent assay was performed as previously described (Uchino et al., 2021). Microtiter plates (Greiner Bio-One, Kremsmünster, Austria, 655061) were coated with 1 µg/mL bovine serum albumin conjugated with RNAPII C-terminal domain peptides with or without phosphorylated amino acids (MAB Institute, Inc.; Figure S6) overnight at 4°C. The plates were washed three times with PBS (Fujifilm Wako Pure Chemical, Osaka, Japan, 048-29805) containing 0.1% Tween-20 (Fujifilm Wako Pure Chemical, 167-11515) (PBST), and each well was incubated with Blocking One P (Nacalai Tesque, 05999-84; 100 µL) for 20 min at RT, washed three times with PBST, and incubated with a 1:3 dilution series of purified RNAPII Ser5ph-mintbody (starting at 300 ng/mL) and IgG antibodies (starting at 30 ng/mL) specific for Ser2ph (CMA602, RRID: AB_2819246) (Stasevich et al., 2014) and Ser5ph (CMA603, RRID: AB_2827955) (Stasevich et al., 2014) in PBST (100 µL) overnight at 4°C. After washing three times with PBST, the plates were incubated with anti-GFP (1:2,000, MBL, 598-7) or anti-mouse IgG (1:10,000, Jackson ImmunoResearch, West Grove, PA, USA, 715-035-150), each conjugated with horseradish peroxidase for 120 min at RT. After washing three times with PBST and incubating for 10 min at RT, the plates were incubated in o-phenylenediamine solution (100 µL; 0.26 mg/mL in 0.1 M sodium citrate, pH 5.0, and 0.01% hydrogen peroxide; Fujifilm Wako Pure Chemical, 158-01671) at RT. The absorbance at 490 nm was measured with a reference wavelength of 600 nm using a Varioskan spectrophotometer (Thermo Fisher Scientific).

### Transcription inhibitor treatment in RNAPII Ser5ph mintbody-expressing cells

Nst-SNAP-Ser5ph cells (Table S1) stably expressing RNAPII Ser5ph mintbody-SNAPtag were established as described above (for details see Establishment of fluorescent protein-expressing cells using the piggyBac system). Nst-SNAP-Ser5ph cells grew normally, suggesting that expression of the RNAPII Ser5ph mintbody did not significantly affect cell growth. Nst-SNAP-Ser5ph cells (7 × 10^4^) were cultured overnight on a laminin-511-coated 8-well chambered cover glass. Nst-SNAP-Ser5ph cells were incubated in 2i medium containing 300 nM SNAP-Cell 647-SiR for 30 min at 37°C and 5% CO_2_ to stain the SNAPtag. After incubation, the cells were washed three times with fresh 2i medium and further incubated for 30 min at 37°C and 5% CO_2_. The medium was replaced with imaging 2i medium containing 15 µM THZ1 (Selleck Chemicals, Houston, TX, USA, S7549), 1 µM flavopiridol (Chemscene LLC, CS-0018), 100 µM DRB, or 0.13% dimethyl sulfoxide, and the cells were further incubated for 1 h before imaging.

Acquired images were filtered with a one-pixel diameter 3D Gaussian Blur filter and subjected to background subtraction with a rolling ball radius of 50 pixels, followed by maximum intensity projection using ImageJ software. Nuclei were selected manually using “Polygon selections” or “Freehand selections” tools with the ROI manager, and the mean intensity and foci number of regions of interest were calculated using the find maxima (*prominence* = 10) and measure functions. The mean intensity of foci in the nucleus was determined by averaging the focus intensities of each cell.

### RF and mintbody imaging and analysis

SNAPtag knock-in cells or mintbody-SNAPtag-expressing mESCs (5 × 10^4^) were plated into a well of a laminin-511-coated 8-well chambered cover glass and cultured overnight at 37°C and 5% CO_2_. The cells were incubated in 2i medium containing 300 nM SNAP-Cell 647-SiR for 30 min at 37°C and 5% CO_2_. After washing the cells three times with 2i medium, they were incubated for another 30 min at 37°C and 5% CO_2_.

The medium was replaced with imaging medium (2i). We focused on the z-position where the mTetR spot was detectable and acquired single-section images in the order of the SNAPtag, MCP, and mTetR channels ten times. To measure the accuracy of the microscope system, images of 0.1 µm fluorescent beads (TetraSpeck Microspheres, Thermo Fisher Scientific, T7279) on an 8-well chamber cover glass in imaging 2i medium were acquired. The images were processed using Gaussian Blur 3D (pixel size = 1) and bleach correction (simple ratio) using ImageJ software. Only SNAPtag images were processed with subtraction background rolling = 50, and all other images were processed with subtraction background rolling = 5. All images were then processed using average intensity projection.

Image analysis of SNAPtag cluster, MCP, and mTetR spots was performed as described by Li et al., 2020 (Li et al., 2020). First, a 19 × 19-pixel region of interest centered on the mTetR spot was selected manually. The coordinates of mTetR and MCP spots were determined by local maxima using trackpy.locate in the Trackpy package.

The spots were then subjected to Gauss fitting using trackpy.refine_leastsq to determine the coordinates at the subpixel level. Next, MCP spots within 390 nm of the mTetR spot with a relative fluorescence intensity greater than two-fold to the average intensity of the 19 × 19-pixel region of interest were classified as the “ON” state, and the remaining spots were classified as “OFF”. SNAPtag clusters were identified by local maxima using trackpy.locate. In addition, the cluster centers were determined at the sub-pixel level by Gauss fitting using trackpy.refine_leastsq. The 2D distances between the nearest SNAP and mTetR coordinates were calculated in each cell.

For time-lapse imaging, five z-sections per stack spanning 0.8 µm (z-step = 200 nm) were acquired at 10 s intervals for 10 min. The images were processed with Gaussian Blur 3D (pixel size = 1) using ImageJ software. Only the SNAPtag images were processed with Subtract background rolling = 50, and all other images were processed with Subtract background rolling = 5. All images were processed using maximum-intensity projection. The coordinates of the RF cluster, mTetR, and MCP spots were determined as described above. As the time-lapse images were noisy to the minimal laser excitation, the locally estimated scatterplot smoothing regression curve was drawn with the degree of smoothing α = 0.2 using R ggplot2. Using these regression curves, the periods spanning at least three consecutive MCP signals less than 2-fold above the background were defined as the OFF state.

### Quantification and statistical analysis

The exact number, *N*, of data points and their representation (such as cells and independent experiments), and statistical tests used are indicated in the respective figure legends and in the results. Statistical tests were performed in R software (The R Project for Statistical Computing, Vienna, Austria). Boxplots with descriptive statistics were created in R software. Boxes indicate the interquartile range (IQR; 25–75% intervals) and median line; whiskers indicate 1.5-fold of the IQR.

### Supplemental item titles and legends

**Fig. S1 Knock-in of STREAMING-tag to *Nanog* and its effects, related to Fig. 1 and 2**. (A) Comparison of amino acid sequences of NeoR-I and NeoR-II. (B) Genomic DNA and coding amino acid sequences of wild-type (WT) around the STREAMING-tag knock-in site in *Nanog*. (C) Effect of STREAMING-tag knock-in into *Nanog* on gene products. (D) STREAMING-tag knock-in site in *Nanog* and distances from transcription start sites to the knock-in site. (E) DNA sequence of the WT allele in NSt cells, in which a deletion of six base pairs was introduced. (F) Single-molecule fluorescence *in situ* hybridization (smFISH) analysis of WT cells. Arrowheads indicate *Nanog* transcriptional spots. Scale bar, 10 µm.

**Fig. S2 Establishment of *Sox2* and *Usp5* STREAMING-tag knock-in cell lines, related to Fig. 2.** (A) Gene structure of mouse *Sox2* before and after STREAMING-tag knock-in. (B) Southern blot analysis of *Sox2* STREAMING-tag knock-in candidate cells. (C) Sequence confirmation of non-knock-in allele in monoallelic knock-in candidate cells. (D) Western blot analysis using SOX2 and GAPDH antibodies in knock-in candidate cells. Clone-10 and Clone-14 cells are biallelic, and Clone-12 and Clone-34 are monoallelic knock-in cells. In this study, we used Clone-12 as a *Sox2* STREAMING-tag knock-in cell. (E) Gene structure of mouse *Usp5* before and after STREAMING-tag knock-in. (F) Southern blot analysis of *Usp5* STREAMING-tag knock-in candidate cells. (G) Sequence confirmation of non-knock-in allele in monoallelic knock-in candidate cells. Multiple peaks were observed in the sequence chromatogram for Clone-16. Therefore, Clone-16 may contain multiple cells with different mutations. (H) Western blot analysis using USP5 and GAPDH antibodies in knock-in candidate cells. In this study, we used Clone-8 as *Usp5* STREAMING-tag knock-in cells.

**Fig. S3 TetR(W43F) fluorescent spot indicates the target gene locus, related to Fig. 3**. DNA-FISH analysis. NSt-derived cells mildly expressing MCP-RFP and TetR(W43F)-mNG were imaged immediately after fixation with 4% paraformaldehyde. The same samples were subjected to DNA-fluorescence *in situ* hybridization using probes against the *Nanog* locus. Arrowheads indicate overlapping spots of MCP, TetR(W43F), and *Nanog* DNA-FISH signals. The dashed lines indicate the nuclei.

**Fig. S4 Establishment of *Pou5f1*, *Wnk1,* and *Flnc* STREAMING-tag knock-in cell lines, related to Fig. 4.** (A) Gene structure of mouse *Pou5f1* before and after STREAMING-tag knock-in. (B) Southern blot analysis of *Pou5f1* STREAMING-tag knock-in candidate cells. Clones marked with red text indicate those that may not have random integration and have the desired cassette inserted in the expected position. We used Clone-2 as a *Pou5f1* STREAMING-tag knock-in cell. (C) Gene structure of mouse *Wnk1* before and after STREAMING-tag knock-in. (D) Southern blot analysis of *Wnk1* STREAMING-tag knock-in candidate cells. Clones marked with red text indicate clones that may not have random integration and have the desired cassette inserted in the expected position. In this study, we used Clone-6 as *Wnk1* STREAMING-tag knock-in cells. (E) Gene structure of mouse *Flnc* before and after STREAMING-tag knock-in. (F) Southern blot analysis of *Flnc* STREAMING-tag knock-in candidate cells. Clones marked with red text indicate those that may not have undergone random integration and have the desired cassette inserted in the expected position. We used Clone-6 as *Wnk1* STREAMING-tag knock-in cells.

**Fig. S5 Distribution of regulatory factor (RF) cluster-mTetR distance in *Nanog* STREAMING-tag knock-in cells, related to Fig. 6 and 7.** (A) 2D distances between mTetR spots and the nearest RF clusters in the ON states. *N*, number of cells analyzed. To investigate chromatic aberrations between the mNG and SNAPtag channels of the microscope used, the spots of the mNG/SNAPtag channels of 0.1 µm fluorescent beads (27 ± 19 nm in 2D) were measured. The distances between all mTetR-RF clusters were significantly larger than the measurement error. *P*-values were determined using Wilcoxon rank sum test. (B) Cumulative density function (CDF) of the data in (A) is shown. (C) Relative mTetR-RF-cluster xy coordinates. Data are classified by transcriptional states. RPB1, *N* = 69 (51); BRD4, *N* = 38 (31); MED19, *N* = 41 (32); MED22, *N* = 43 (40); Ser5ph, *N* = 74 (63); Ser2ph, *N* = 69 (51), at the ON state (OFF state).

**Fig. S6 Establishment of a mintbody that recognizes RNAPII Ser5ph, related to Fig. 7**. (A) Schematic representation of the mintbody that binds to phosphorylated Ser5 on the C-terminal domain of RNA polymerase II (RNAPII). (B) Example images of mutant scFv-sfGFP stably expressed in HeLa cells. The image acquisition and contrast adjustment settings were the same, enabling comparison of mintbody localization between samples. Scale bar, 10 µm. (C) Nuclear-to-cytoplasm fluorescence intensity ratios of 44B12 and mutants. HeLa cells expressing 44B12-sfGFP and mutants were established. After confocal image acquisition, the nuclear-to-cytoplasmic fluorescence intensity ratios were measured (*N* = 20). (D) Comparison of amino acid sequences of RNAPII Ser5ph (44B12) and H4K20me1 (15F11) scFv. The framework region (FR), complementarity determining region (CDR), and linker region are shown. Mutation sites in 44B12 are indicated in red letters. The final construct, named as RNAPII Ser5ph-mintbody, contains T26A and M83L substitutions. (E–F) His-tagged RNAPII Ser5ph-mintbody was expressed in *Escherichia coli*, purified using an Ni-column, treated with enterokinase to remove the His-tag, and further purified. (E) Purified proteins were separated on an SDS-polyacrylamide gel and stained with Coomassie blue. The positions of the size marker are shown on the left. (F) Enzyme-linked immunosorbent assay plates coated with synthetic peptides conjugated with bovine serum albumin were incubated with a dilution series of purified RNAPII Ser5ph-mintbody and control antibodies specific for RNAPII Ser2ph (CMA602) and RNAPII Ser5ph (CMA603). After incubation with peroxidase-conjugated anti-GFP (for RNAPII Ser5ph-mintbody) or anti-mouse IgG (for monoclonal antibody) and then with o-phenylenediamine, the absorbance was measured at 490 nm. RNAPII Ser5ph-mintbody reacted with peptides containing Ser5ph. (G) Images showing Nst-SNAP-Ser5ph cells stably expressing RNAPII Ser5ph mintbody-SNAPtag cultured with transcriptional inhibitors. Nst-SNAP-Ser5ph cells were labeled with SNAP-Cell 647-SiR and cultured for 1 h either with 15 µM THZ1, 1 µM flavopiridol (FP), 100 µM 5,6-dichloro-1-β-d-ribofuranosylbenzimidazole (DRB) or dimethyl sulfoxide (DMSO) as a vehicle. Scale bar, 5 µm. (H–J) Boxplots with all data points showing mean intensity of RNAPII Ser5ph-mintbody in the nucleus (H), number of RNAPII Ser5ph-mintbody foci in the nucleus (I), and mean intensity of RNAPII Ser5ph-mintbody foci in each cell (J). *P*-values were determined using Wilcoxon rank sum test. DMSO, *N* = 39; THZ1, *N* = 55; FP, *N* = 38; DRB, *N* = 54.

**Fig. S7 Method for establishing cells that moderately express mTetR-mNG, MCP-RFP, and mintbody-SNAP-tag.** mTetR-mNG binds specifically to the TetO sequence. By inserting TetO repeats into a specific genomic region and expressing mTetR-mNG, the region can be visualized. This procedure can also be used for MS2 repeat/MCP-RFP and RNAPII Ser2ph or Ser5ph/mintbody. However, the expression levels of fluorescent protein fusion proteins should be carefully monitored. (A) Visualization of the TetO repeat requires an appropriate amount of mTetR-mNG expression. If the amount of mTetR-mNG is extremely low, the number of mTetR-mNG molecules that bind to the target region will be extremely low, making it difficult to recognize the mTetR spot using fluorescence microscopy (left panel). If there is excess mTetR-mNG, the number of mTetR-mNG molecules bound to the target region will be high. However, if the concentration of unbound mTetR-mNG in the cell nucleus is higher than the local concentration of mTetR-mNG bound to TetO repeats, recognition of the mTetR spot is difficult (right panel). If the appropriate amount of mTetR-mNG is expressed, the local concentration of mTetR-mNG bound to TetO repeats will be higher than that of unbound mTetR-mNG in the cell nucleus, allowing for recognition of the mTetR spot (middle panel). Therefore, it is necessary to select cells expressing an appropriate amount of mTetR-mNG. The same procedure can be used for MS2 repeat/MCP-RFP and RNAPII Ser2ph or Ser5ph mintbody. (B) Fluorescence-activated cell sorting (FACS) gating strategy for cells expressing appropriate amounts of mTetR-mNG, MCP-RFP, and mintbody-SNAPtag in mouse embryonic stem cells (mESCs). First, we prepared cells stably expressing mTetR-mNG, MCP-RFP, and mintbody-SNAPtag using the piggyBac system in mESCs (see STAR Methods). From these cell populations, cells expressing appropriate amounts of mTetR-mNG, MCP-RFP, and mintbody-SNAPtag were sorted using FACS. Shown is an example of an FACS gating strategy in mTetR-mNG-, MCP-RFP-, and RNAPII Ser2ph mintbody-SNAPtag-expressing cells. Immediately before FACS, the cells were treated with SNAP-Cell 647-SiR and stained with SNAPtag. The P8 gate is a cell population that does not express mNG or RFP. Approximately, 300 cells from the P16 gate were sorted using FACS and plated into 6 cm dishes. The resulting colonies were passaged on an 8-well chambered cover glass with #1.5 glass (Cellvis, catalog number: C8-1.5HN) for microscopic observation, and cells expressing the appropriate amount of mNG/RFP/SNAPtag were selected using microscopy.

## Methods S1: Detailed step-by-step protocols related to STAR Methods

### Construction of STREAMING-tag knock-in vectors

#### Materials

- Left-arm-F primer (see below)
- Left-arm-R primer (see below)
- Right-arm-F primer (see below)
- Right-arm-R primer (see below)
- KOD One polymerase (TOYOBO, KMM-101X5) or another high-fidelity DNA polymerase
- Agarose S (Nippon Gene, 313-90231)
- TBE buffer (Takara Bio, T9122)
- Zymoclean™ Gel DNA Recovery Kits (Zymo Research, D4002)
- BsaI-HFv2 (NEB, R3733S)
- SacI-HF (NEB, R3156S)
- KpnI-HF (NEB, R3142S)
- pBSKΔBS-SD_MCS_SA(NBSI)-24xMS2-sirius-96xopto-TetO-Int-3xFLAG-NeoR-GS4 (Addgene ID # 177265).
- pBSKΔB (Addgene ID # 177268)
- NEBuilder HiFi DNA Assembly kit (NEB, E2621L)
- One Shot™ Stbl3™ Chemically Competent (Thermo Fisher Scientific, C737303)
- LB medium (Nacalai Tesque, 20068-75)
- Ampicillin (Nacalai Tesque, 02739-74)
- GenElute HP Plasmid Miniprep Kit (Sigma-Aldrich, NA0160-1KT)

#### Protocols

1. Determine the knock-in site of the STREAMING-tag cassette based on the genome sequence of the target species. Ideally, the knock-in site should be near the transcription start site (TSS) and translated by all transcripts. Considering the following points, it is necessary to knock-in at appropriate sites in the coding region near the TSS. First, it has been reported that the length of the first exon may affect transcriptional activity (Bieberstein et al., 2012). As the splice donor is just below the 5′ side of the STREAMING-tag cassette, the exon length may be altered by the STREAMING-tag knock-in. Moreover, because of the N-end rule, knock-in of the STREAMING-tag immediately after the initiation codon should be avoided (Tasaki et al., 2012). Thus, it is considered to knock-in the STREAMING-tag at least three amino acids downstream from the starting methionine. In addition, some proteins may contain a domain near the N-terminus, which is important for protein function. As STREAMING-tag knock-in results in insertion of an amino acid sequence, QGCF (Fig. S1C), the knock-in position should be determined on a case-by-case basis. Because the 5′ and 3′ ends of the STREAMING-tag both encode amino acids (Fig. S1C), the knock-in site should be set between the codon sequences to be in-frame. In addition, there must be a highly specific CRISPR target sequence that overlaps the desired knock-in position. For mice and humans, highly specific CRISPR target sequences can be easily identified using the UCSC genome browser (http://genome.ucsc.edu). In addition, the insertion site should be designed to be located 10–20 bases from the spacer sequence of the CRISPR target sequence, which will prevent the CRISPR from being re-cut after knock-in. Once the insertion position was determined, the primer sets were designed. The following primer set was used to target the *Sox2* locus: Left-arm-F (5′-CTATAGGGCGAATTGGGTACGAGCGCAGTGCCGCGGATGAGCGC-3′), Left-arm-R (5′-GATCGAATACTTATCGCTTA<u>CCTG</u>GCCGGTCGCCGCCGCCGTGGCGTT-3′), Right-arm-F (5′-GGTCTGGTTGCAAGCAATTG<u>CTTC</u>GGCAACCAGAAGAACAGCCCGGAC-3′), and Right-arm-R (5′-AGGGAACAAAAGCTGGAGCTAATGGGCCTTAAAAATACCAGCGG-3′).
2. Red letters indicate homologous sequence sites between fragments that will be used when assembling with NEBuilder, and underlined green letters indicate additional sequences for codon frame adjustment. Depending on the target sequence, the black part can be modified appropriately. We designed the left and right arms to be approximately 1 kbp in length. NCBI Primer-BLAST (https://www.ncbi.nlm.nih.gov/tools/primer-blast/) was used for primer design. These oligo DNAs can be synthesized by an appropriate company.
3. Amplify the left arm and right arm separately by PCR using high-fidelity thermostable DNA polymerase (such as KOD One polymerase, TOYOBO, Osaka, Japan) using the prepared oligo DNA and genomic DNA of the target species as templates.
4. Resolve the PCR products on 1% agarose gel in 0.5× TBE buffer.
5. Cut out approximately 1 kbp fragments of the left and right arm fragments from the agarose gels.
6. Purify the DNA from the gel slabs using Zymoclean™ Gel DNA Recovery Kits and elute the DNA with 10 µL of ddH2O. Measure the DNA concentration using a Nanodrop to ensure correct stoichiometry in the following steps. If not used, immediately freeze and store the DNA frozen until needed.
7. Treat the pBSKΔBS-SD_MCS_SA(NBSI)-24xMS2-sirius-96xopto-TetO-Int-3xFLAG-NeoR-GS4 (Table S2) with BsaI. Resolve the digestion product on a 1% agarose gel in 0.5×TBE buffer.
8. Cut out the ∼5.5 kbp fragment of pBSKΔBS-SD_MCS_SA(NBSI)-24xMS2-sirius-96xopto-TetO-Int-3xFLAG-NeoR-GS4 BsaI digest from the agarose gels.
9. Purify the DNA from the gel slabs using Zymoclean™ Gel DNA Recovery Kits and elute the DNA with 10 µL of ddH2O. This fragment is named as the STREAMING-tag fragment. Measure the DNA concentration with a Nanodrop to ensure correct stoichiometry in the following steps. If not used, immediately freeze and store the DNA until needed.
10. Treat pBSKΔB (Table S2) with SacI and KpnI. Resolve the digest on a 1% agarose gel in 0.5×TBE buffer.
11. Cut out the ∼3 kbp fragment of pBSKΔB SacI/KpnI digest from the agarose gels.
12. Purify the DNA from the gel slabs using Zymoclean™ Gel DNA Recovery Kits and elute the DNA with 10 µL of ddH2O. This fragment is named as the backbone fragment. Measure the DNA concentration using a Nanodrop to ensure the correct stoichiometry in the following steps. If not used, immediately freeze and store the sample until required.
13. Use the NEBuilder HiFi DNA Assembly kit to assemble these fragments together.
14. Prepare the fragments in advance by diluting them with ddH2O to the following concentrations.

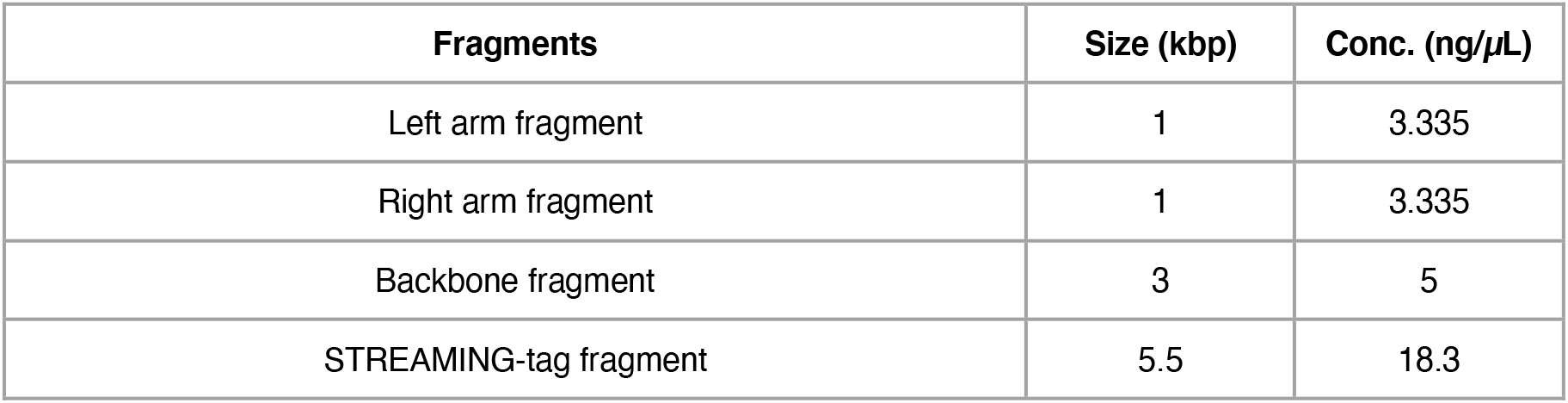
15. Mix the samples in the following combinations: Prepare a reaction solution with only the backbone added as a negative control.

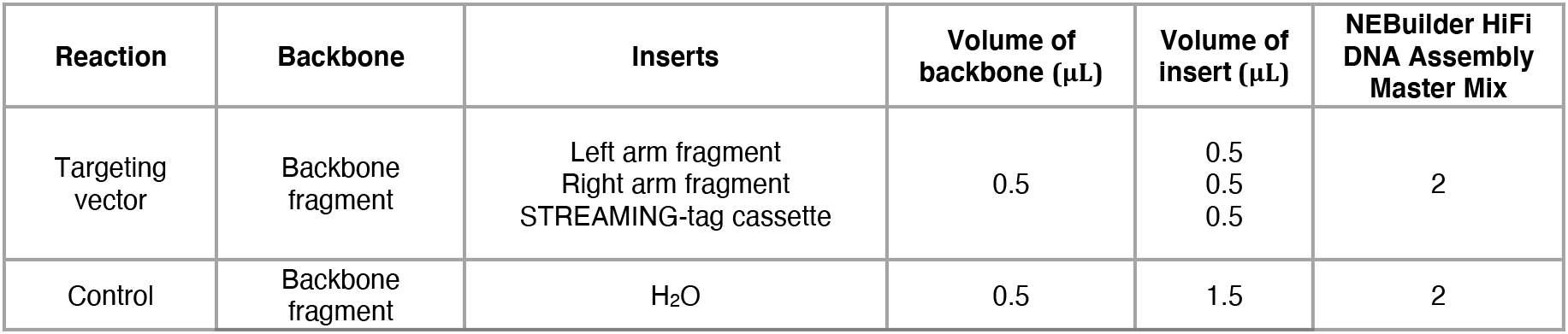
16. Incubate the sample in a thermocycler at 50°C for 60 min, and then place the samples on ice.
17. Transform the targeting vector and control reactions into 50 µL of Stbl3 chemically competent cells as per the manufacturer’s protocol.
18. Spread *Escherichia coli* onto LB agarose medium containing ampicillin and incubate at 30°C overnight.
19. If the number of colonies does not exceed twice that of the control, most plasmids were not successfully created. In this case, it is recommended that the samples be reconditioned.
20. Seed the resulting colonies individually into 3 mL of LB medium containing ampicillin and incubate the cultures overnight at 32°C with shaking.
21. Collect the plasmid with the GenElute HP Plasmid Miniprep Kit.
22. Treat the recovered plasmid with appropriate restriction enzymes and verify the expected band pattern.
23. As the homology arm region amplified using PCR and junction sites in NEBuilder HiFi-assembled plasmids are prone to mutation, these regions should be confirmed using Sanger sequencing using appropriate oligo DNA.
24. The final plasmid should be purified using an appropriate method to achieve transfection-grade purity.

#### Construction of the CRISPR vector Materials

- eSpCas9-EF (Addgene ID # 177267)
- BbsI-HF (NEB, R3539S)
- Agarose S (Nippon gene, 313-90231)
- TBE buffer (Takara Bio, T9122)
- Zymoclean™ Gel DNA Recovery Kits (Zymo Research, D4002)
- Sense oligo (see below)
- Antisense oligo (see below)
- T4 Polynucleotide Kinase (PNK) (NEB, M0201S)
- ATP (NEB, P0756S)
- Ligation-Convenience Kit (Nippon gene, 319-05961)
- One Shot™ Stbl3™ Chemically Competent (Thermo Fisher Scientific, C737303)
- LB medium (Nacalai Tesque, 20068-75)
- Ampicillin (Nacalai Tesque, 02739-74)
- GenElute HP Plasmid Miniprep Kit (Sigma-Aldrich, NA0160-1KT)
- LKO1-5 primer (5′-GACTATCATATGCTTACCGT-3′)

#### Protocols

1. Treat the eSpCas9-EF plasmid (Table S2) with BbsI, and resolve the digest on a 1% agarose gel in 0.5× TBE buffer.
2. Cut out the ∼8.5 kbp fragment of eSpCas9-EF BbsI digest from the agarose gels.
3. Purify the DNA from the gel slabs using Zymoclean™ Gel DNA Recovery Kits, and elute the DNA with 10 µL of ddH2O. This fragment is named as the backbone fragment. Measure the DNA concentration with a Nanodrop to ensure the correct stoichiometry in the following steps. If not used, immediately store the fragment frozen until needed.
4. Order the following oligo DNA containing the spacer sequence corresponding to the CRISPR target sequence determined above (5′-GCTGTTCTTCTGGTTGCCGC<u>CGG</u> -3′ for Sox2; underlined part indicates the PAM sequence). Sense oligo: 5′-CACC<u>G</u>GCTGTTCTTCTGGTTGCCGC-3′, antisense oligo: 5′-AAACGCGGCAACCAGAAGAACAGC<u>C</u>-3′. The red letters are additional sequences for cloning, and underlined parts indicate an additional G base at the 5′ end of the spacer sequence. This additional G base is inserted because it is required for transcription initiation by the U6 promoter. If the 5′ end of the spacer sequence is G, this additional G base does not need to be inserted.
5. To perform phosphorylation and annealing of oligos, prepare the following reaction solutions in PCR tubes. Finally, add the relevant oligonucleotides.

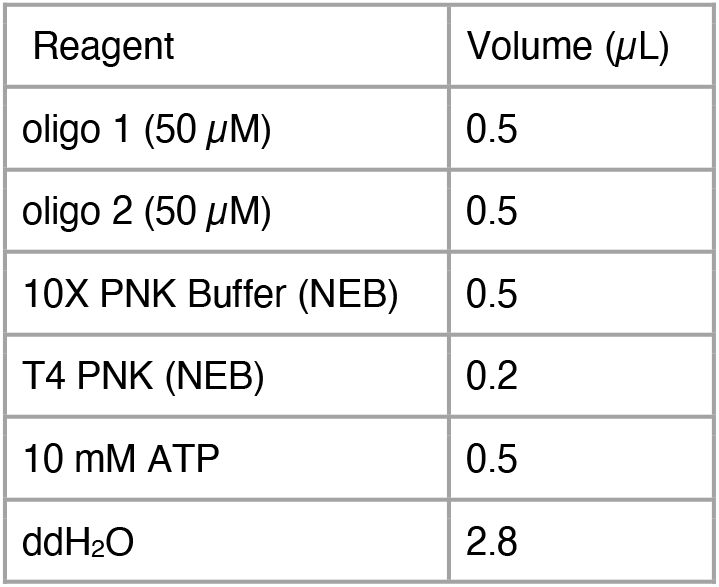
6. Mix the reaction solution well by tapping or pipetting, and collect the sample at the bottom of the tube by centrifugation.
7. Perform the following reaction in a thermal cycler to anneal and phosphorylate the oligos: 37°C for 30 min, 95°C for 5 min, and then ramp down to 25°C at 5°C/min.
8. After the reaction, collect the sample at the bottom of the tube by centrifugation and place the sample on ice. This fragment is named as the spacer fragment.
9. Remove the Ligation-Convenience Kit from the freezer, thaw if frozen, and place on ice.
10. Prepare the samples as follows: backbone should be pre-diluted as appropriate.

| Reagent | Volume (µL) |
| --- | --- |
| oligo 1 (50 µM) | 0.5 |
| oligo 2 (50 µM) | 0.5 |
| 10X PNK Buffer (NEB) | 0.5 |
| T4 PNK (NEB) | 0.2 |
| 10 mM ATP | 0.5 |
| ddH<sub>2</sub>O | 2.8 |

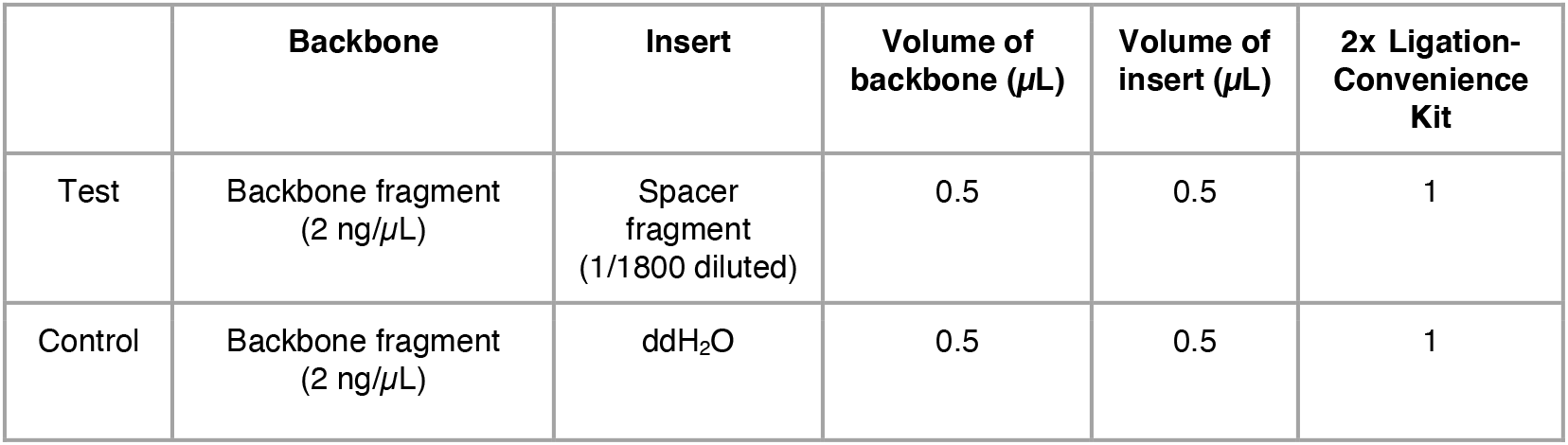
11. Mix the reaction contents well by pipetting and allow to ligate at 16°C for 30 min.
12. Transform the sample into Stbl3 chemical competent cells according to the manufacturer’s protocol.
13. Spread *Escherichia coli* onto LB agarose medium containing ampicillin, and incubate the cells at 37°C overnight.
14. If the number of colonies does not exceed twice that of the control, it is likely that most plasmids were not successfully prepared. In this case, it is recommended that the samples be reconditioned.
15. Place the resulting colonies individually into 3 mL of LB medium containing ampicillin, and incubate the cultures overnight at 37°C with shaking.
16. Purify the plasmid using the GenElute HP Plasmid Miniprep Kit.
17. Treat the recovered plasmid with appropriate restriction enzymes, and confirm that the expected band pattern is obtained.
18. As the homology arm region amplified using PCR and junction sites in NEBuilder HiFi-assembled plasmid are prone to mutations, these regions should be verified using Sanger sequencing using the LKO1-5 primer (5′-GACTATCATATGCTTACCGT-3′).
19. Purify the final plasmid using an appropriate method to achieve transfection-grade purity.

### Establishment of STREAMING-tag knock-in cells

Here, we present an example of generating STREAMING-tag knock-in mESCs. When using different cell types, it is necessary to optimize the transfection method and G418 treatment concentration depending on the cell type.

#### Materials

- C57BL6J mESC cell line (Bruce 4 C57BL/6J, male, EMD Millipore, Billerica, MA, USA)
- Targeting vector (see above)
- CRISPR vector (see above)
- pKLV-PGKpuro2ABFP (Addgene # 122372)
- 2i medium (Dulbecco’s modified Eagle’s medium [DMEM]; 15% fetal bovine serum [FBS]; 0.1 mM β-mercaptoethanol; 1× MEM nonessential amino acids; 2 mM L-alanyl-L-glutamine solution; 1,000 U/mL leukemia inhibitory factor [LIF]; 20 µg/mL gentamicin; 3 µM CHIR99021; and 1 µM PD0325901)
- Opti-MEM reduced serum medium (Life Technologies, Carlsbad, CA, USA; 11058021)
- Lipofectamine 3000 (Life Technologies, L3000015)
- Puromycin (Wako, 160-23151)
- G418 (Nacalai Tesque, 16512-81)

#### Protocols

1. Day 0

1. Plate mESCs (5 × 10^5^) into each well of a 12-well plate, and after 1 h, mix the following transfection reagents: 2 µg of targeting vector (e.g., pTV-Nanog_2-5prime-1000-24MS96T-3F_NeoR), 700 ng of CRISPR vector (e.g., eSpCas9-EF-5Nanog_2), and 300 ng of pKLV-PGKpuro2ABFP.
2. To each of these, add 62.5 µL of reduced serum Opti-MEM and 2.5 µL of P3000 reagent.
3. In a separate tube, add 62.5 µL of reduced serum Opti-MEM and 4.5 µL of Lipofectamine 3000 per reaction mixture, and mix well.
4. Mix these reagents in equal volumes and incubate for 15 min at room temperature. Add the complex to wells containing cells, and incubate overnight.
2. Day 1

Replace the medium with fresh 2i medium containing 1 µg/mL puromycin.
3. Day 2

Replace the medium with fresh 2i medium.
4. Day 3

Transfer all cells to gelatin-coated 10 cm dishes.
5. Day 5

Replace the medium with 8 mL of 2i medium containing 200 µg/mL G418.
6. Day 7

Replace the medium with 8 mL of 2i medium containing 200 µg/mL G418.
7. Day 9

After another 48 h, pick up 24 colonies.

Genomic DNA is extracted from a portion of each colony, and knock-in candidate colonies are selected using genomic PCR. In addition, after expanding the candidate colonies, genomic DNA is extracted from a portion of each colony, and the rest is cryopreserved. Knock-in and random integration of vectors are confirmed using Southern blotting.

### Establishment of fluorescent protein-expressing cells using the piggyBac system

Here, we present an example of mESCs. When using different cell types, it is necessary to optimize the transfection method and G418 treatment concentration depending on the cell type.

#### Materials

- STREAMING-tag knock-in mESC lines
- 2i medium (Dulbecco’s modified Eagle’s medium [DMEM]; 15% fetal bovine serum [FBS]; 0.1 mM β-mercaptoethanol; 1×MEM nonessential amino acids; 2 mM L-alanyl-L-glutamine solution; 1,000 U/mL leukemia inhibitory factor [LIF]; 20 µg/mL gentamicin; 3 µM CHIR99021; and 1 µM PD0325901)
- pCAG hyPBase (Ochiai et al., 2015)
- pLR5-CAG-TetR_W43F-3xmNG (Addgene #174880)
- pLR5-CAG-hMCP-mScarlet-I-NLS (Addgene #174878)
- SNAPtag expression vector (e.g., pLR5-CAG-NLS-SNAP, Addgene #174879)
- Opti-MEM reduced serum medium (Life Technologies, Gaithersburg, MD, USA; 11058021)
- Lipofectamine 3000 (Life Technologies, L3000015)
- SNAP-Cell 647-SiR (New England Biolabs, S9102S)

#### Protocols

1. Day 0

1. Plate STREAMING-tag knock-in mESC lines (2.5 × 10^5^) to each well of a 24-well plate; after 1 h, mix the transfection reagents as described below.
2. In a tube, mix 50 ng pCAG hyPBase, 75 ng pLR5-CAG-TetR_W43F-3xmNG, 275 ng pLR5-CAG-hMCP-mScarlet-I-NLS, and 100 ng of SNAPtag expression vector (pLR5-CAG-NLS-SNAP).
3. To each of these, add 25 µL of reduced serum Opti-MEM and 1 µL of P3000 reagent.
4. In a separate tube, add 25 µL of reduced serum Opti-MEM and 1.8 µL of Lipofectamine 3000 per reaction mixture, and mix well.
5. Mix these reagents in equal volumes and incubate at room temperature for 15 min.
2. Day 1

Replace the medium with 2i medium.
3. Day 2

Replace the medium with 2i medium.
4. Day 5

Passage the cells to 12-well plates.
5. Day 6

1. Incubate the cells in 2i medium containing 300 nM SNAP-Cell 647-SiR for 30 min at 37 °C and 5% CO_2_ for another 30 min.
2. Collect the cells by trypsin treatment, and cells moderately expressing mTetR-mNG, MCP-RFP, and SNAPtag should be sorted using a BD FACSAria III cell sorter (BD Biosciences, Franklin Lakes, NJ, USA) and seeded onto gelatin-coated 6 cm dishes. The cells should show a mild level of fluorescence expression; if the expression level is too high, it will be difficult to detect the clusters (Fig. S7).
6. Day 8

Replace the medium with 2i medium, and change the medium once every 2 days.
7. Day 11

Observe cells expressing moderate levels of fluorescent protein under a fluorescence microscope, and use these cells for further experiments.

**Movie S1. Dynamics of MS2 coat protein (MCP) and mTetR spots in NSt-NLS-SNAP cells.** NSt-NLS-SNAP cells were treated with SNAP-Cell 647-SiR immediately before imaging. The movie shows a maximum intensity projection-processed time-lapse image of NSt-NLS-SNAP cells captured at 15 s intervals. Scale bar, 5 µm.

**Table S1. Cell lines used in this study**

**Table S2. Plasmids used in this study**

**Table S3. Southern blot probes and primers used to generate the probes**

