## Supplementary Figs and Methods for "STREAMING-tag system reveals spatiotemporal relationships between transcriptional regulatory factors and transcriptional activity"

**Fig. S7 Method for establishing cells that moderately express mTetR-mNG, MCP-RFP, and mintbody-SNAP-tag.** mTetR-mNG binds specifically to the TetO sequence. By inserting TetO repeats into a specific genomic region and expressing mTetR-mNG, the region can be visualized. This procedure can also be used for MS2 repeat/MCP-RFP and RNAPII Ser2ph or Ser5ph/mintbody. However, the expression levels of fluorescent protein fusion proteins should be carefully monitored. (A) Visualization of the TetO repeat requires an appropriate amount of mTetR-mNG expression. If the amount of mTetR-mNG is extremely low, the number of mTetR-mNG molecules that bind to the target region will be extremely low, making it difficult to recognize the mTetR spot using fluorescence microscopy (left panel). If there is excess mTetR-mNG, the number of mTetR-mNG molecules bound to the target region will be high. However, if the concentration of unbound mTetR-mNG in the cell nucleus is higher than the local concentration of mTetR-mNG bound to TetO repeats, recognition of the mTetR spot is difficult (right panel). If the appropriate amount of mTetR-mNG is expressed, the local concentration of mTetR-mNG bound to TetO repeats will be higher than that of unbound mTetR-mNG in the cell nucleus, allowing for recognition of the mTetR spot (middle panel). Therefore, it is necessary to select cells expressing an appropriate amount of mTetR-mNG. The same procedure can be used for MS2 repeat/MCP-RFP and RNAPII Ser2ph or Ser5ph mintbody. (B) Fluorescence-activated cell sorting (FACS) gating strategy for cells expressing appropriate amounts of mTetR-mNG, MCP-RFP, and mintbody-SNAPtag in mouse embryonic stem cells (mESCs). First, we prepared cells stably expressing mTetR-mNG, MCP-RFP, and mintbody-SNAPtag using the piggyBac system in mESCs (see STAR Methods**)**. From these cell populations, cells expressing appropriate amounts of mTetR-mNG, MCP-RFP, and mintbody-SNAPtag were sorted using FACS. Shown is an example of an FACS gating strategy in mTetR-mNG-, MCP-RFP-, and RNAPII Ser2ph mintbody-SNAPtag-expressing cells. Immediately before FACS, the cells were treated with SNAP-Cell 647-SiR and stained with SNAPtag. The P8 gate is a cell population that does not express mNG or RFP. Approximately, 300 cells from the P16 gate were sorted using FACS and plated into 6 cm dishes. The resulting colonies were passaged on an 8-well chambered cover glass with #1.5 glass (Cellvis, catalog number: C8-1.5HN) for microscopic observation, and cells expressing the appropriate amount of mNG/RFP/SNAPtag were selected using microscopy.

### Fig. S1

**A**

```
35.1% identity in 251 residues overlap; Score: 367.0; Gap frequency: 2.8%

NeoR-I        12 LYGYKWARDNVGQSGATIYRLYGKPDAPELFLKHGKGSVANDVTDEMVRLNWL-TEFMPL
NeoR-II       25 LFGYDWAQQTIGCSDAAVFRLSAQ-GRPVLFVKTDLSGALNELQDEAARLSWLATTGVPC
                 * ** **    * * *   **      * ** *       *   **  ** ** *   *

NeoR-I        71 PTIKHFIRTPDDAWLLTTAIPGKTAFQVLEEYPDSGENIVDALAVFLRRLHSIPVCNCPF
NeoR-II       84 AAVLDVVTEAGRDWLLLGEVPGQ---DLLSSHLAPAEK-VSIMADAMRRLHTLDPATCPF
                              ***    **      *       *  *   *   ****      ***

NeoR-I       131 NSDRVFRLAQAQSRMNNGLVDASDFDDERNGWPVEQVWKEMHKLLPFSPDSVVTHGDFSL
NeoR-II      140 DHQAKHRIERARTRMEAGLVDQDDLDEEHQGLAPAELFARLKARMPDGDDLVVTHGDACL
                       *   *  **  ****  * * *  *              *   * ******  *

NeoR-I       191 DNLIFDEGKLIGCIDVGRVGIADRYQDLAILWNCLGE-FSPSLQKRLFQKYGIDNPDMNK
NeoR-II      200 PNIMVENGRFSGFIDCGRLGVADRYQDIALATRDIAEELGGEWADRFLVLYGIAAPDSQR
                  *     *   * ** ** * ****** *       *        *    ***  **

NeoR-I       250 LQFHLMLDEFF
NeoR-II      260 IAFYRLLDEFF
                   *   *****
```

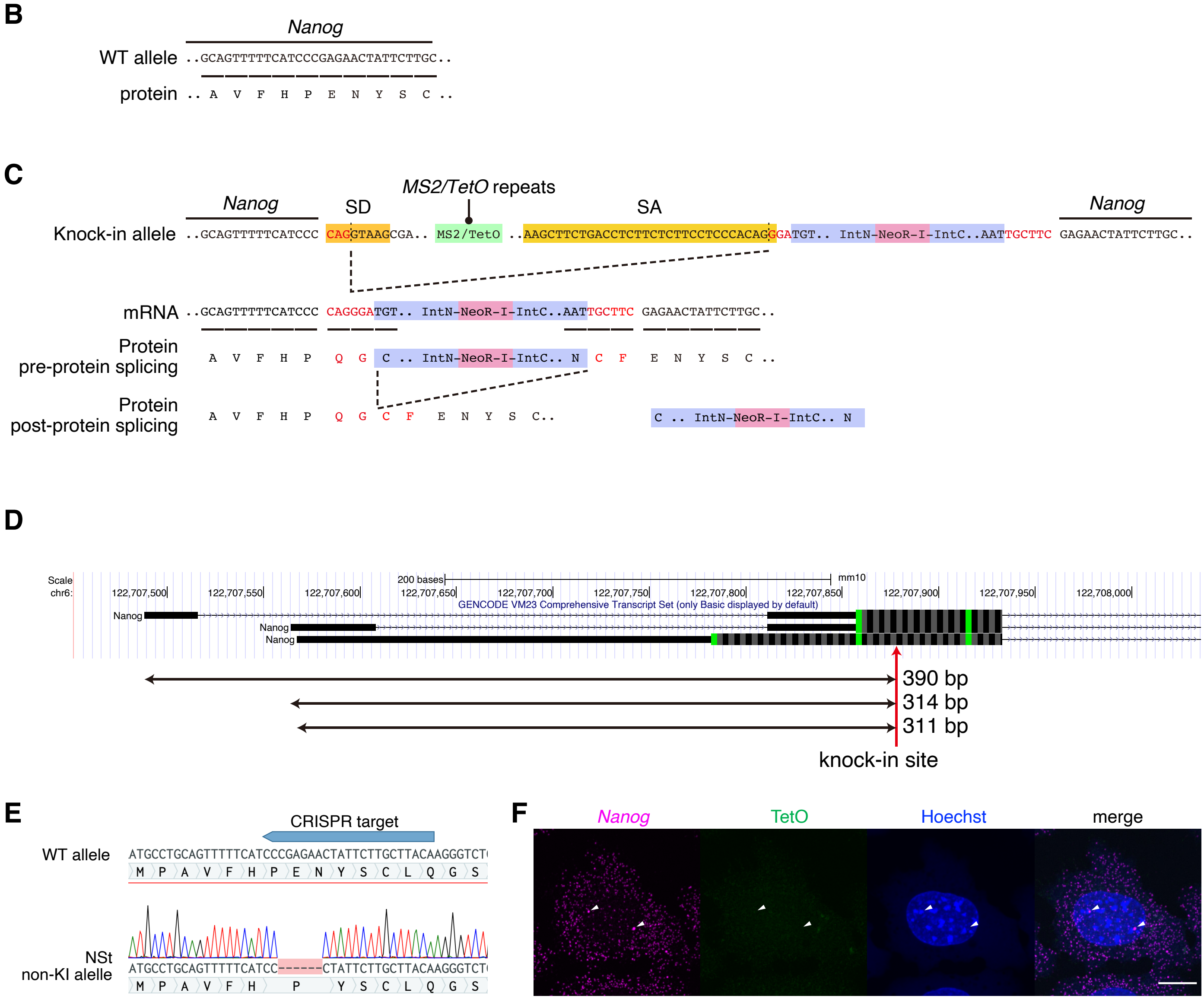

#### Fig. S2

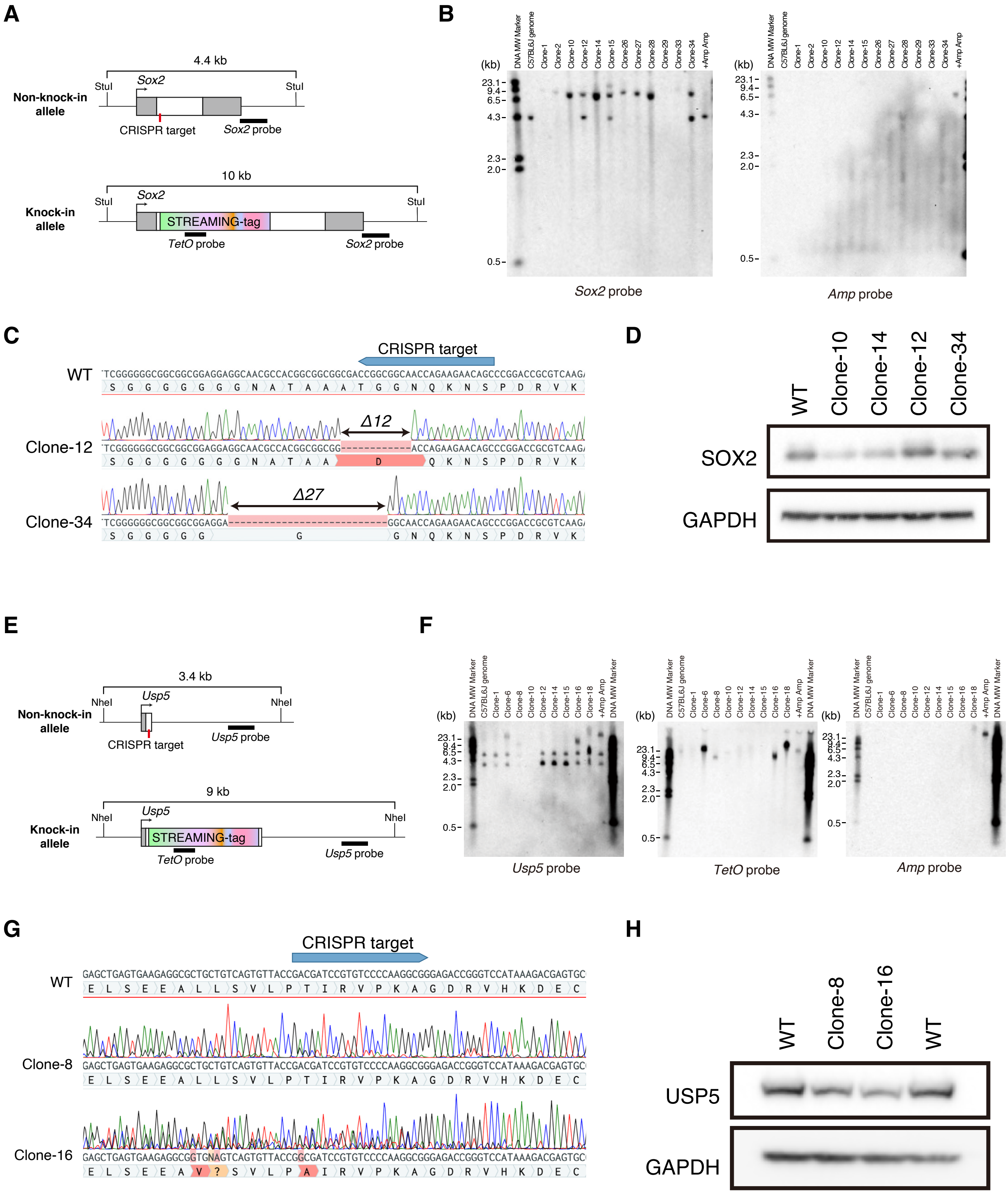
A
4.4 kb
Non-knock-in allele
StuI
Sox2
CRISPR target
Sox2 probe
10 kb
Knock-in allele
STREAMING-tag
TetO probe
B
Sox2 probe
Amp probe
C
CRISPR target
WT
Clone-12
Δ12
Clone-34
Δ27
D
WT
Clone-10
Clone-14
Clone-12
Clone-34
SOX2
GAPDH
E
3.4 kb
Non-knock-in allele
NheI
Usp5
CRISPR target
Usp5 probe
9 kb
Knock-in allele
STREAMING-tag
TetO probe
F
Usp5 probe
TetO probe
Amp probe
G
CRISPR target
WT
Clone-8
Clone-16
H
WT
Clone-8
Clone-16
USP5
GAPDH

**Fig. S3**

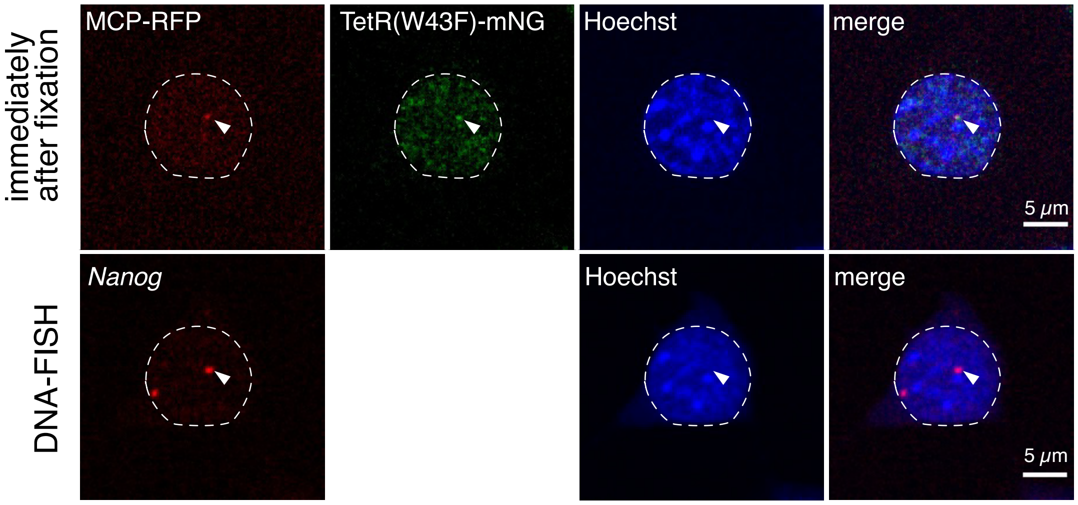
MCP-RFP
TetR(W43F)-mNG
Hoechst
merge
immediately after fixation
5 μm
*Nanog*
Hoechst
merge
DNA-FISH
5 μm

Fig. S4

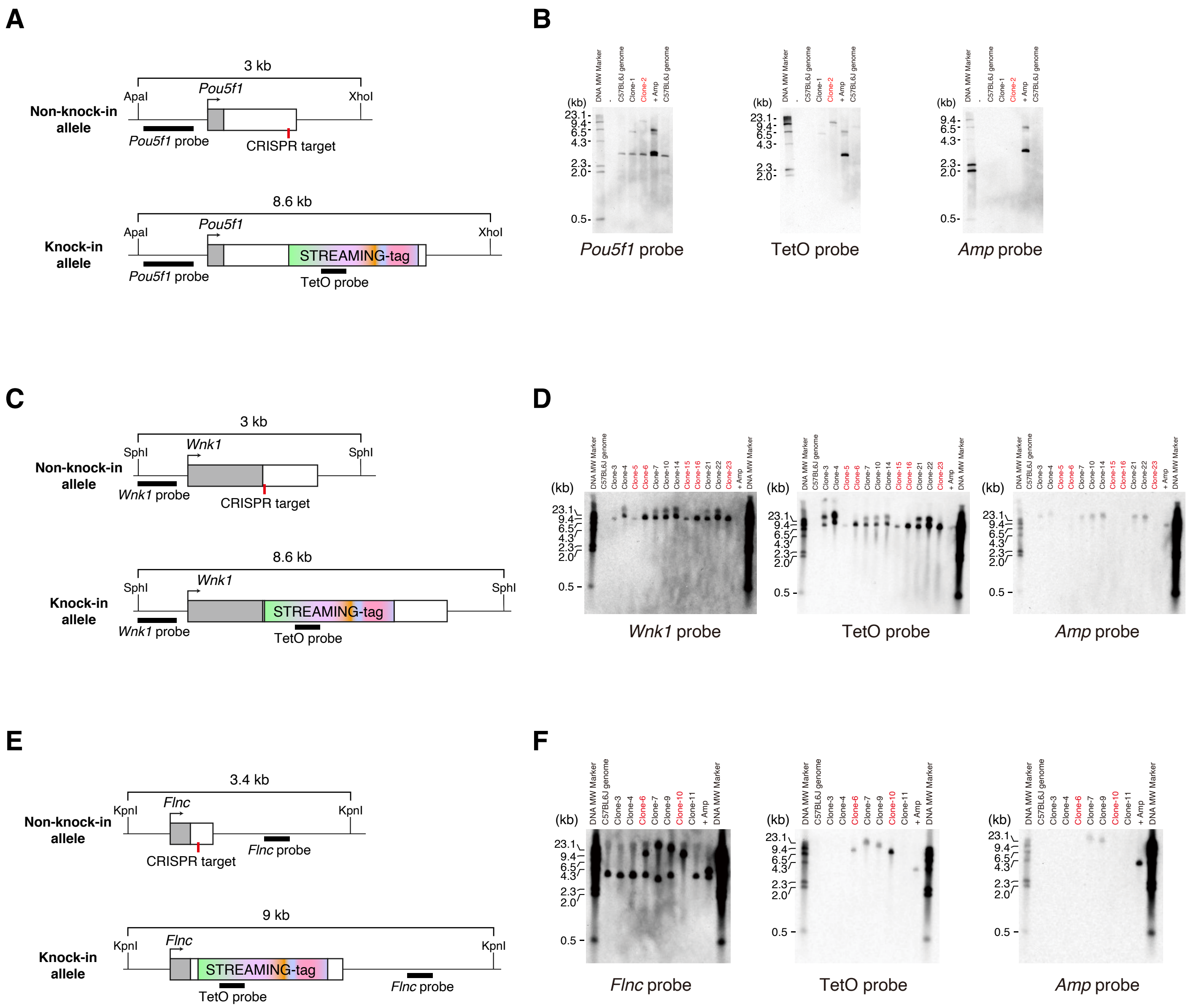
A
3 kb
Non-knock-in allele
ApaI
Pou5f1
XhoI
Pou5f1 probe
CRISPR target
8.6 kb
Knock-in allele
ApaI
Pou5f1
XhoI
STREAMING-tag
Pou5f1 probe
TetO probe
B
(kb)
23.1
9.4
6.5
4.3
2.3
2.0
0.5
DNA MW Marker
C57BL6J genome
Clone-1
Clone-2
+ Amp
Pou5f1 probe
TetO probe
Amp probe
C
3 kb
Non-knock-in allele
SphI
Wnk1
SphI
Wnk1 probe
CRISPR target
8.6 kb
Knock-in allele
SphI
Wnk1
SphI
STREAMING-tag
Wnk1 probe
TetO probe
D
(kb)
23.1
9.4
6.5
4.3
2.3
2.0
0.5
DNA MW Marker
C57BL6J genome
Clone-3
Clone-4
Clone-5
Clone-6
Clone-7
Clone-10
Clone-14
Clone-15
Clone-16
Clone-21
Clone-22
Clone-23
+ Amp
Wnk1 probe
TetO probe
Amp probe
E
3.4 kb
Non-knock-in allele
KpnI
Flnc
KpnI
CRISPR target
Flnc probe
9 kb
Knock-in allele
KpnI
Flnc
KpnI
STREAMING-tag
TetO probe
Flnc probe
F
(kb)
23.1
9.4
6.5
4.3
2.3
2.0
0.5
DNA MW Marker
C57BL6J genome
Clone-3
Clone-4
Clone-6
Clone-7
Clone-9
Clone-10
Clone-11
+ Amp
Flnc probe
TetO probe
Amp probe

**Fig. S5**

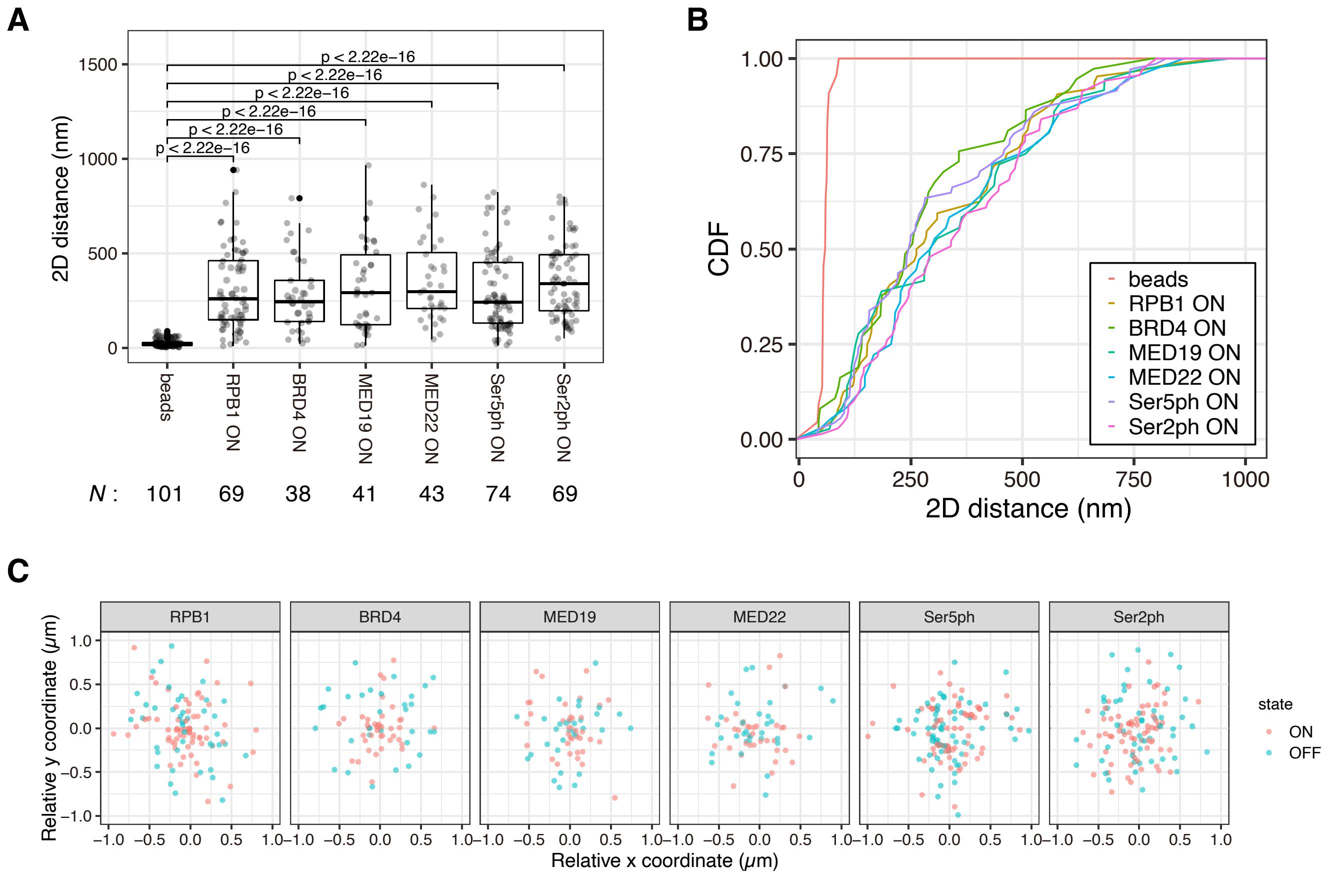
A
2D distance (nm)
1500
1000
500
0
p < 2.22e−16
p < 2.22e−16
p < 2.22e−16
p < 2.22e−16
p < 2.22e−16
p < 2.22e−16
beads
RPB1 ON
BRD4 ON
MED19 ON
MED22 ON
Ser5ph ON
Ser2ph ON
N : 101 69 38 41 43 74 69
B
CDF
1.00
0.75
0.50
0.25
0.00
0
250
500
750
1000
2D distance (nm)
beads
RPB1 ON
BRD4 ON
MED19 ON
MED22 ON
Ser5ph ON
Ser2ph ON
C
RPB1
BRD4
MED19
MED22
Ser5ph
Ser2ph
Relative y coordinate (μm)
Relative x coordinate (μm)
state
ON
OFF

**Fig. S6**

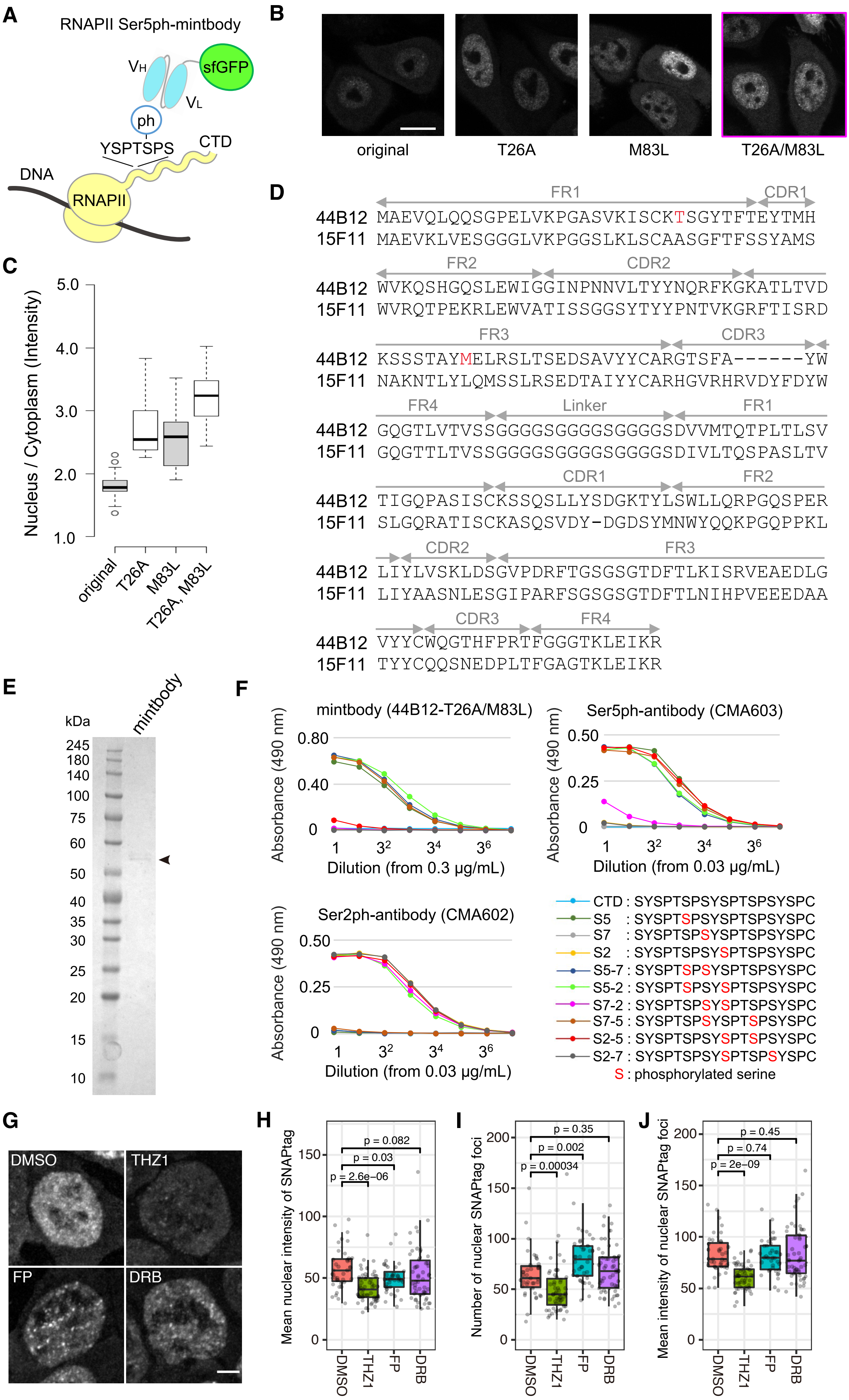
A
RNAPII Ser5ph-mintbody
VH
sfGFP
VL
ph
YSPTSPS
CTD
DNA
RNAPII
B
original
T26A
M83L
T26A/M83L
C
Nucleus / Cytoplasm (Intensity)
5.0
4.0
3.0
2.0
1.0
original
T26A
M83L
T26A, M83L
D
FR1
CDR1
44B12 MAEVQLQQSGPELVKPGASVKISCKTSGYTFTEYTMH
15F11 MAEVKLVESGGGLVKPGGSLKLSCAASGFTFSSYAMS
FR2
CDR2
44B12 WVKQSHGQSLEWIGGINPNNVLTYYNQRFKGKATLTVD
15F11 WVRQTPEKRLEWVATISSGGSYTYYPNTVKGRFTISRD
FR3
CDR3
44B12 KSSSTAYMELRSLTSEDSAVYYCARGTSFA------YW
15F11 NAKNTLYLQMSSLRSEDTAIYYCARHGVRHRVDYFDYW
FR4
Linker
FR1
44B12 GQGTLVTVSSGGGGSGGGGSGGGGSDVVMTQTPLTLSV
15F11 GQGTTLTVSSGGGGSGGGGSGGGGSDIVLTQSPASLTV
CDR1
FR2
44B12 TIGQPASISCKSSQSLLYSDGKTYLSWLLQRPGQSPER
15F11 SLGQRATISCKASQSVDY-DGDSYMNWYQQKPGQPPKL
CDR2
FR3
44B12 LIYLVSKLDSGVPDRFTGSGSGTDFTLKISRVEAEDLG
15F11 LIYAASNLESGIPARFSGSGSGTDFTLNIHPVEEEDAA
CDR3
FR4
44B12 VYYCWQGTHFPRTFGGGTKLEIKR
15F11 TYYCQQSNEDPLTFGAGTKLEIKR
E
kDa
mintbody
245
180
140
100
75
60
50
40
35
30
25
20
15
10
F
mintbody (44B12-T26A/M83L)
Absorbance (490 nm)
0.80
0.40
0
1
3²
3⁴
3⁶
Dilution (from 0.3 µg/mL)
Ser5ph-antibody (CMA603)
0.50
0.25
Dilution (from 0.03 µg/mL)
Ser2ph-antibody (CMA602)
CTD : SYSPTSPSYSPTSPSYSPC
S5 : SYSPTSPSYSPTSPSYSPC
S7 : SYSPTSPSYSPTSPSYSPC
S2 : SYSPTSPSYSPTSPSYSPC
S5-7 : SYSPTSPSYSPTSPSYSPC
S5-2 : SYSPTSPSYSPTSPSYSPC
S7-2 : SYSPTSPSYSPTSPSYSPC
S7-5 : SYSPTSPSYSPTSPSYSPC
S2-5 : SYSPTSPSYSPTSPSYSPC
S2-7 : SYSPTSPSYSPTSPSYSPC
S : phosphorylated serine
G
DMSO
THZ1
FP
DRB
H
Mean nuclear intensity of SNAPtag
p = 0.082
p = 0.03
p = 2.6e−06
150
100
50
0
DMSO
THZ1
FP
DRB
I
Number of nuclear SNAPtag foci
p = 0.35
p = 0.002
p = 0.00034
200
150
100
50
0
J
Mean intensity of nuclear SNAPtag foci
p = 0.45
p = 0.74
p = 2e−09
200
150
100
50
0

**Fig. S7**

**A**

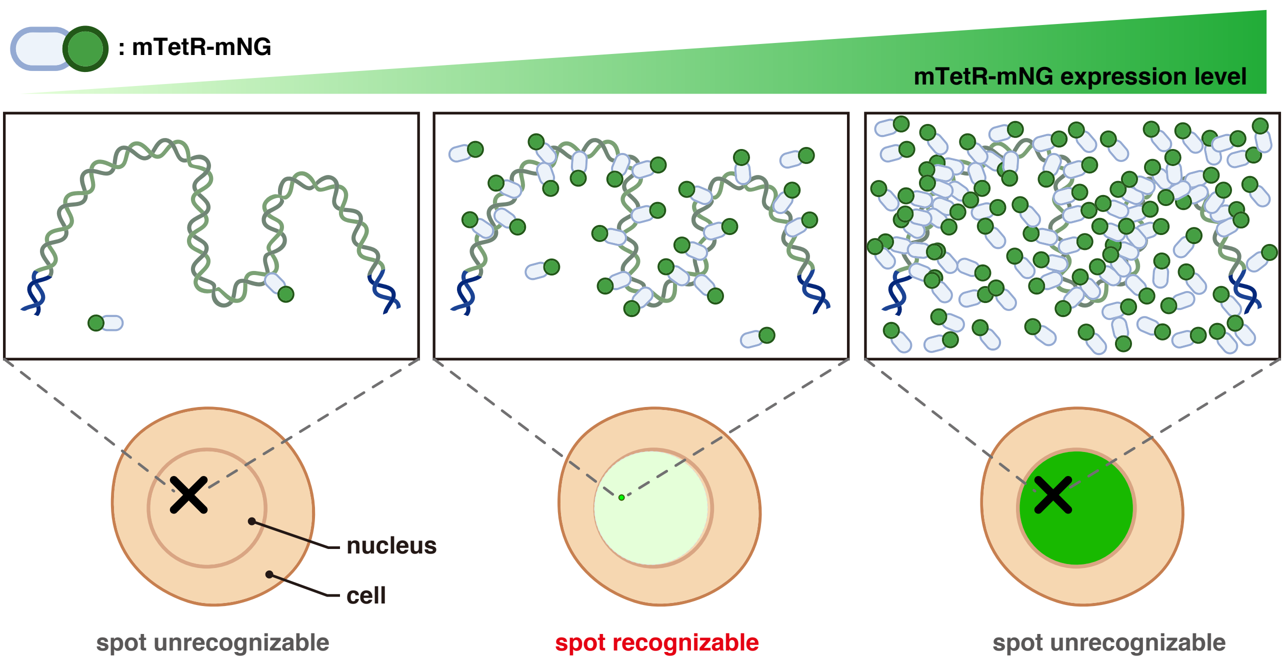

: mTetR-mNG
mTetR-mNG expression level
nucleus
cell
spot unrecognizable
spot recognizable
spot unrecognizable


**B**

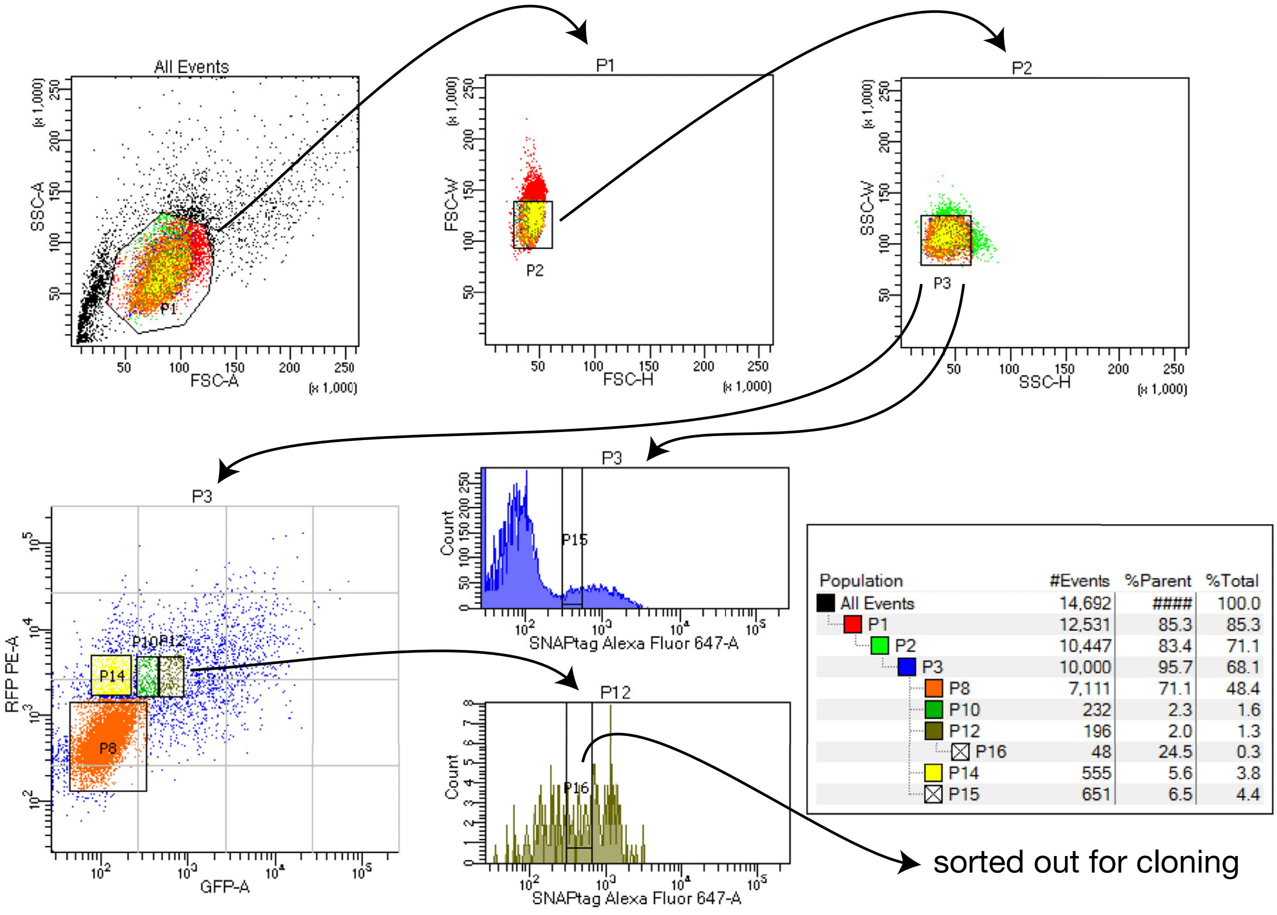

All Events
SSC-A
FSC-A
(x 1,000)
P1
FSC-W
FSC-H
P2
SSC-W
SSC-H
P3
RFP PE-A
GFP-A
P10
P12
P14
P8
Count
SNAPtag Alexa Fluor 647-A
P15
P16
Population #Events %Parent %Total
All Events 14,692 #### 100.0
P1 12,531 85.3 85.3
P2 10,447 83.4 71.1
P3 10,000 95.7 68.1
P8 7,111 71.1 48.4
P10 232 2.3 1.6
P12 196 2.0 1.3
P16 48 24.5 0.3
P14 555 5.6 3.8
P15 651 6.5 4.4
sorted out for cloning

| **Fragments** | **Size (kbp)** | **Conc. (ng/µL)** |
|---|---|---|
| Left arm fragment | 1 | 3.335 |
| Right arm fragment | 1 | 3.335 |
| Backbone fragment | 3 | 5 |
| STREAMING-tag fragment | 5.5 | 18.3 |

15. Mix the samples in the following combinations: Prepare a reaction solution with only the backbone added as a negative control.

| Reaction | Backbone | Inserts | Volume of backbone (μL) | Volume of insert (μL) | NEBuilder HiFi DNA Assembly Master Mix |
|---|---|---|---|---|---|
| Targeting vector | Backbone fragment | Left arm fragment<br>Right arm fragment<br>STREAMING-tag cassette | 0.5 | 0.5<br>0.5<br>0.5 | 2 |
| Control | Backbone fragment | $H_2O$ | 0.5 | 1.5 | 2 |

| | **Backbone** | **Insert** | **Volume of backbone (μL)** | **Volume of insert (μL)** | **2x Ligation-Convenience Kit** |
|---|---|---|---|---|---|
| Test | Backbone fragment (2 ng/μL) | Spacer fragment (1/1800 diluted) | 0.5 | 0.5 | 1 |
| Control | Backbone fragment (2 ng/μL) | $ddH_2O$ | 0.5 | 0.5 | 1 |
